# Global distribution of isoprenoid quinones across Bacteria

**DOI:** 10.1101/2025.09.17.676790

**Authors:** Sophie-Carole Chobert, Suraj Kanwar, Olivier Lerouxel, Nelle Varoquaux, Julie Michaud, Ludovic Pelosi, Fabien Pierrel, Sophie S. Abby

## Abstract

Isoprenoid quinones represent a class of redox lipids involved in many critical cellular functions, including ATP synthesis through electron transport chains. They thus occupy a pivotal role in the bioenergetics of all three domains of life. The diversity of quinone types observed across microbial taxa has long supported their use as chemotaxonomic markers in microbial systematics. More recently, variations in quinone repertoires have been linked to metabolic adaptations and a novel quinone was discovered. Despite a revived interest in the role of quinones, a unified perspective on the distribution of quinones in Bacteria is currently lacking. In this study, quinone biosynthetic pathways were systematically annotated in 26,264 high quality genomes of bacterial species, and specific information on quinones produced by over 6,000 bacterial species was extracted by text mining the abstracts of thousands of articles. The results were mapped onto a phylogenetic tree, providing the most comprehensive overview of quinone distribution in Bacteria to date. This enabled us to highlight the surprisingly dynamic evolutionary history of the two menaquinone-producing pathways. Moreover, the identification and experimental validation of a deeply branching ubiquinone pathway in *Desulfobacterota* represents the first occurrence of such a pathway outside the *Pseudomonadota* and provides insights into the nature of the ancestral UQ pathway. The updated compendium of bacterial quinones is a valuable resource to facilitate the prediction of quinone structural features from genomic data, to establish correlations between quinone structures and cellular traits, and to explore the evolution of quinone repertoires in connection with the diversification of microbial metabolisms.

## Introduction

ATP production by ATP synthase is nearly ubiquitous and relies on chemical gradients generated by electron transfer chains (ETCs). Isoprenoid quinones are redox lipids that transfer electrons between proteins of the ETCs, making them central to bioenergetics as key components in both respiration and photosynthesis. Hence, they are virtually found in any organism except for purely fermentative species and methanogens. Besides this well-established role in bioenergetics, isoprenoid quinones (henceforth referred to as “quinones”) are also involved in several important cellular processes, including haem and pyrimidine synthesis, and disulfide bond formation [1–3]. They act as protectants against oxidative and osmotic stresses. For instance, high menaquinone levels in membrane composition of halophilic archaea (>70%) were proposed to contribute to tolerance towards high salt concentrations and oxidative stress [4]. Further, quinone content in membranes was proposed to be involved in adaptation to low growth temperature in *Listeria monocytogenes* [5]. However, the link between quinone content and structure, and e.g. membrane fluidity, is still unclear. Quinones are also involved in regulating the metabolic shifts between aerobiosis and anaerobiosis [6], and they function as extracellular growth factors in syntrophic gut bacteria [7].

Isoprenoid quinones are found in the three domains of life. Menaquinone (MK) is widespread in bacteria and archaea. Plastoquinone (PQ) and phylloquinone (PhK) are key for oxygenic photosynthesis in cyanobacteria and plants, and ubiquinone (UQ) is found in the bacterial phylum *Pseudomonadota* and in eukaryotes, where it is often called “coenzyme Q” [1]. Isoprenoid quinones are composed of a quinone ring conjugated to a polyisoprenyl tail, which is highly hydrophobic and anchors quinones in membranes (Figs 1 and S1). The ring of the most commonly found quinones is either of the benzoquinone (UQ, PQ) or the naphthoquinone (MK, PhK) type. The chemical groups (methyl, methoxy, amino) decorating the quinone rings modulate the redox potential. For example, replacing a methoxy group with an amino group converts ubiquinone (UQ) to rhodoquinone (RQ) and lowers the redox potential by ∼160 mV. This changes the reactivity towards the fumarate/succinate redox couple, with succinate oxidation coupled to UQ and fumarate reduction coupled to RQ [8,9]. The redox potential of quinones therefore determines the respiratory substrates that can be used by the ETC, and consequently the enzymes they function with [10,11].

**Fig 1.**
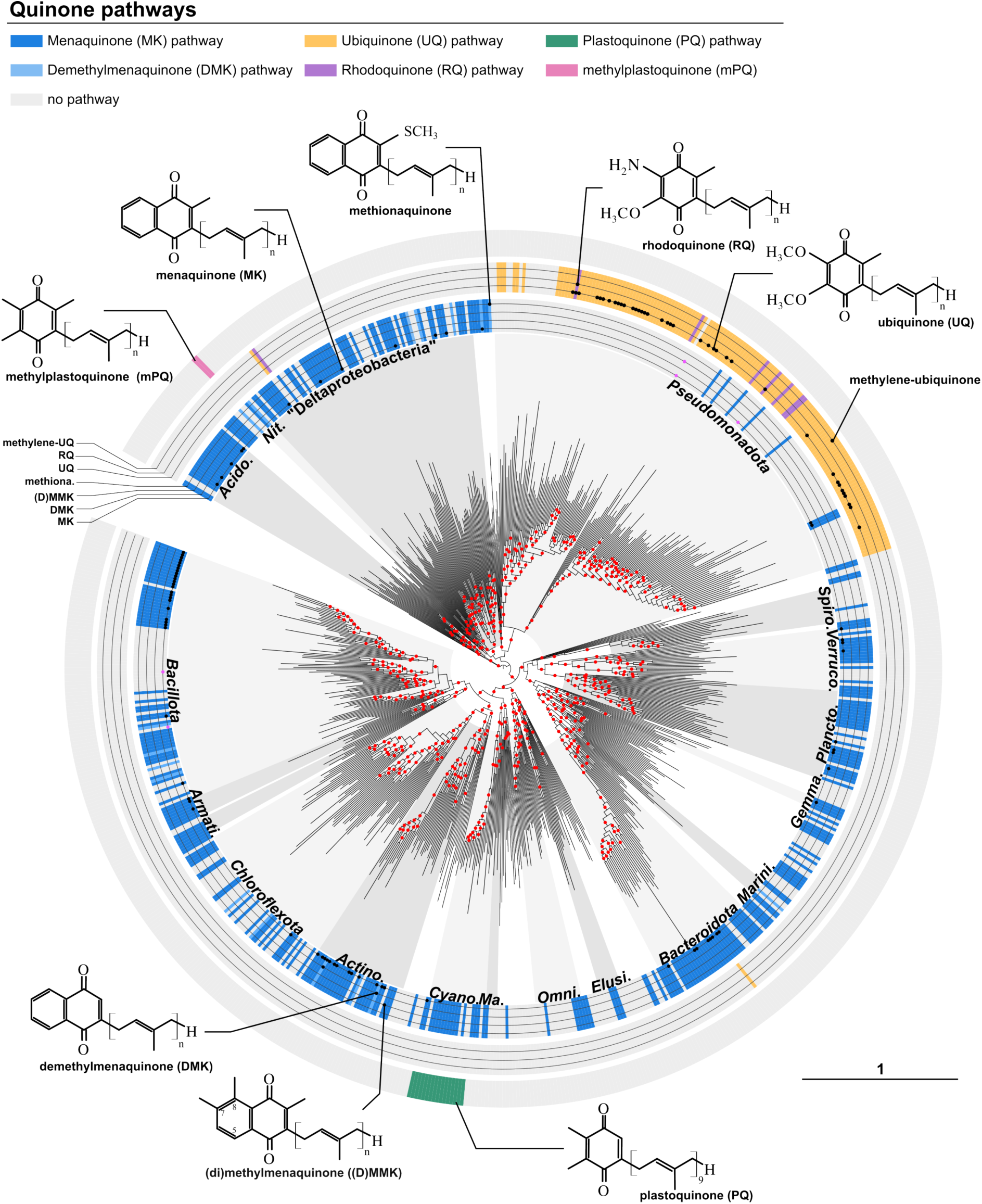
Phylogenetic tree of bacteria showing the presence of quinones from bioinformatic pre-dictions and literature mining. The maximum-likelihood species tree includes 751 high-quality genomes and metagenome-assembled genomes and was obtained from the concatenation of 120 GTDB markers. The tree was rooted to separate the Gracilicutes from the Terrabacteria as previously proposed [47,48]. The following selected phyla (GTDB) are labeled on the tree: *Pseudomonadota, Spirochaetota, Verrucomicrobiota, Planctomycetota, Gemmatimonadota, Marinisomatota, Bacteroidota, Elusimicrobiotoba, Omnitrophota, Margulisbacteria, Cyanobacteriota, Actinomycetota, Chloroflexota, Armatimonadota, Bacillota, Acidobacteriota, Nitrospirota,* and “Deltaproteobacteria” *(Desulfobacterota,*

The length of the polyisoprenyl tail of quinones varies between organisms. For example, UQ and MK in *Escherichia coli* have tails composed of 8 isoprenyl units and are therefore abbreviated UQ_8_ and MK_8_, whereas humans synthesize UQ_10_, *Pseudomonas aeruginosa* has UQ_9_, and *Bacillus subtilis* has MK_7_. The length of the tail is determined by polyprenyl diphosphate synthases but remains difficult to predict from protein sequences and structures [12]. The polyprenyl diphosphate synthases often do not exhibit exquisite selectivity, resulting in a major tail length *n* but also minor amounts of *n-1* and *n+1* isoprenologs, as documented for example in *Acetobacter aceti*, which contains UQ_9_ as the major quinone, and UQ_8_ and UQ_10_ as minor isoprenologs [13]. The tail varies not only in length but also in degree of saturation. Indeed, the C=C double bond of one or more isoprenyl units may be reduced, resulting in partially or fully saturated tails, which will decrease the flexibility of membranes [4]. Gram-positive bacteria often contain MK with one or two saturations, designated MK_n_(H_2_) and MK_n_(H_4_), respectively [13], while several archaeal species belonging to the *Nitrososphaeria* (formerly Thaumarchaeota) or the *Desulfurococcales* contain a fully saturated MK_6_ (designated MK_6:0_, according to a different nomenclature) [14]. However, given that the reductases responsible for the saturations are rarely known [15] and that chain length prediction from polyprenyl diphosphate synthases sequences are inaccurate [12] obtaining reliable quinone tail information from large-scale genomic analyses is currently impossible.

The biosynthesis of isoprenoid quinones depends on multi-step pathways, where the conjugation of the pol4yprenyl tail occurs either on a preformed quinone ring (MK, PhK) or on a ring precursor requiring further modifications (UQ, PQ, RQ). Most steps in the biosynthesis of the commonly found quinones (UQ, PQ, RQ, MK, PhK) are now characterized [8,16–23]. This opens the possibility of scanning genomes for genes encoding quinone biosynthetic enzymes and inferring the production of a given quinone by a species based on the presence of the corresponding biosynthesis pathway. We have used this strategy to study the distribution of UQ, RQ, and MK in *Pseudomonadota*, which enabled the investigation of their evolutionary history and their contribution to metabolic adaptation to environments with varying oxygen levels [24]. Interestingly, a novel quinone, methyl-PQ (mPQ), has recently been discovered in some species belonging to the bacterial phylum *Nitrospirota* [25,26]. Several steps of mPQ biosynthesis are catalyzed by enzymes homologous to those involved in the UQ, PQ, and MK pathways. Two pathways exist for MK biosynthesis, the so-called Men and futalosine pathways [16,17]. Phylogenetic analyses suggest that MK is likely ancestral, with UQ, PQ, and mPQ subsequently diverging from a shared biosynthetic origin [11,25,27,28] Besides these major quinones, undermethylated or overmethylated forms of MK are also known, such as demethylmenaquinone (DMK) or (di)methylmenaquinone ((D)MMK) [29]. Rare and peculiar quinones, whose biosynthetic pathways are not yet known, have also been found in certain organisms [14,30–32] (S1 Fig).

Pioneering systematic surveys of bacterial quinones in the 1960s and 1970s showed that closely related bacteria often share the same type of quinone with the same chain length and saturation [13]. This provided the basis for using isoprenoid quinones as chemotaxonomic markers to facilitate the classification of prokaryotic taxa [33]. As a result, it became common practice to biochemically characterize the quinones synthesized by each newly isolated bacterial species and to report this information in the abstract of the associated publication. In the 1990s, quinone profiling was even used to assess the composition and seasonal variation of bacterial communities in natural environments [34,35]. While information about the type(s) of isoprenoid quinones found in some specific bacterial taxa was collected in the book series *<u>The Prokaryotes</u>* [36–41], the distribution of isoprenoid quinone structural types in bacteria was last compiled by Collins and Jones in 1981 [13]. Although quinones play important roles in cellular, metabolic and ecological processes, our understanding of their diversity and taxonomic distribution remains incomplete, particularly given our expanding knowledge of the tree of life provided by new genome and metagenome sequences. In this study, we present an updated compendium of bacterial quinones. This was compiled by mining the literature for biochemically characterized quinones and their associated species, and by conducting a genomic survey in which known quinone biosynthetic pathways were annotated across the current bacterial diversity. By mapping our data on a phylogenetic tree of *Bacteria*, we provide a global overview of quinone distribution while shedding light and opening perspectives on the evolution of cells, energy metabolism, and quinone biosynthetic pathways.

## Results

### Genomic annotation of quinone pathways in the tree of *Bacteria*

We downloaded 26,264 high-quality representative bacterial genomes from the Genome Taxonomy Database (GTDB) and used a similarity search approach to annotate the biosynthetic genes of the known quinone pathways: UQ, RQ, MK (Men and futalosine pathways), and PQ. To assign complete pathways, a quorum of genes was applied to each pathway as previously described (see Methods and [24]). As the selection of GTDB representative genomes is based on genomic diversity [42], our approach provides an overview of quinone pathway distribution across the currently known bacterial diversity (Figs 1 and S1).

This overview confirmed the wide distribution and dominance of MK, which is believed to be ancient [27,43] (Fig 1 and S1 Table). MK was indeed found in 51% of the taxonomic orders included in this study. The respective distribution of the Men and futalosine pathways is discussed in a dedicated section. As expected, the PQ pathway was found only in the oxygenic phototrophs (*Cyanobacteriia*) of the phylum *Cyanobacteriota.* The UQ pathway was found in most genomes (98%) of the *Pseudomonadota* phylum, which is very diverse as it gathers 159 orders according to the GTDB (91% containing UQ). Unexpectedly, UQ presence was inferred in three lineages outside of the *Pseudomonadota*: the GWC2-55-46 class from *Desulfobacterota*, and two genomes from *Bacillota* and *Bacteroidota* species (GCA_003513005.1 and GCA_020635905.1). The former case is very well supported and is further discussed below. The latter two cases are likely to correspond to annotation errors as not all essential genes of the UQ pathway are present, nor are they closely localized in the two genomes, whereas they usually form several compact genetic loci in *Pseudomonadota* [24]. In bacteria, RQ is produced by the RquA protein, which substitutes a methoxy group of UQ by an amine group [44]. As expected, RQ production was inferred from certain genomes of *Pseudomonadota* lineages that produce UQ, but also in one genome from the GWC2-55-46 class, which harbors the genetic potential for UQ production. Interestingly, as the GTDB dataset covered a wide diversity of *Pseudomonadota* species, our analysis revealed a wider taxonomic distribution of RQ than previously observed (from 8 orders in Chobert *et al.* [24] to 24 orders in this study, see Table S1).

From this overview, it was striking to see that many lineages seem devoid of quinone pathways. These include mostly *Bacillota* and *Chloroflexota*, as well as scattered lineages across the bacterial tree of life [45]. Several lineages are known to contain fermentative species, and, as such, may not require respiratory quinones [45,46]. However, the recent discovery of mPQ, a new quinone in a late emerging lineage *of Nitrospirota* [25,26], could suggest that other quinones are yet to be discovered and may populate various parts of the bacterial tree.

*Myxoccocota, Bdellovibrionota* and relatives). Quinones predictions are depicted in surrounding colored circles: MK and DMK are shown in shades of blue, with vivid blue for MK, lighter blue for DMK, and very light blue when neither is predicted. UQ and RQ are represented by yellow and purple, respectively, with very light yellow indicating neither. PQ predictions and mPQ from experimentally tested species [24] are depicted in green for PQ, pink for mPQ, and very light green when PQ is not predicted. Quinones obtained from abstract mining are indicated on the corresponding sub-rings (MK, DMK, (D)MMK, methionaquinone, UQ, RQ, and methylene-ubiquinone), with black dots when the results of genomic annotations are congruent with the abstract mining, or with pink dots when they are not congruent (see S1 Text). UFBoot branch support values are depicted by red dots when over 95% and the tree scale is expressed in substitutions per site. A labeled version of this tree is presented in S2 Fig.

### Harvesting quinone data from the literature

Since quinones are routinely characterized when new bacterial species are described, we hypothesized that information about experimentally validated quinones would be abundant in the literature. New species are primarily described in four microbiology journals for which 29,268 abstracts were downloaded from the Pubmed database (see Methods), gathering articles from 1948 to 2023. We designed a text mining approach to identify quinone mentions and the species described for which we subsequently obtained taxonomic information and genome sequences, when available (see Methods). This resulted in the extraction of quinone content for 6,382 different species from 5,910 articles, covering 23 different phyla and 145 orders (S2 Table). Most quinone types were found several times in this dataset, except for PQ produced by photosynthetic *Cyanobacteriota*, since *Cyanobacteriota* species are reported in different journals where describing quinone content is uncommon. *Cyanobacteriota* are known to possess a limited range of quinones, including PQ_9_ and either PhK or MK_4_ [21,22].

The collected dataset was highly biased towards certain phyla, as 41% consisted of *Pseudomonadota* species (subset of formerly named Proteobacteria), 24% of *Actinomycetota* (formerly Actinomycetes), and 20% of *Bacteroidota* (formerly Bacteroidetes). The results of the text mining and quinone pathway genomic annotations could be compared for 4,824 species and were overwhelmingly consistent (∼98%) (Fig 1). Most cases of discordance correspond to 39 species for which experimental characterizations reported MK rather than DMK, as inferred from our genomic prediction based the absence of a *menG/ubiE* methyltransfer-ase-encoding gene [24]. Additionally, MK was measured in four species, none of which had a MK pathway in their genome (see S1 Text). Finally, the text mining approach specifically retrieved several rare quinones, for which genomic inference is either complicated by unreliable detection of the corresponding genes (*menK*, *menK2*, and *mqnK* for MMK and DMMK biosynthesis) or simply impossible because the corresponding biosynthetic enzymes are unknown (for methylene-ubiquinone, methionaquinone, or ω-cyclic menaquinone). Overall, the data obtained through the text mining approach provides strong support for the genomic annotation of the quinone biosynthesis pathways, while also highlighting gaps in our knowledge.

### The two MK pathways show a mutually exclusive and highly intricate phylogenetic distribution

MK can be produced via the Men pathway or the more recently discovered futalosine pathway [17,20]. A previous evolutionary study examined the distribution of these two pathways and concluded that the futalosine pathway was more widely distributed and transmitted mostly vertically, whereas the Men pathway appeared to be restricted to fewer lineages and more frequently horizontally transferred [27]. By annotating the MK production pathways in a more diverse set of bacterial genomes, we were able to investigate the distribution and dynamics of the two pathways in more detail (Figs 1 and S2, S1 Table). We confirmed that, although the futalosine pathway is found in fewer genomes than the Men pathway (4,217 *vs* 8,991, S1 Table), it is distributed across a more diverse range of bacterial orders (293 *vs* 208 for Men, Fig 2A). The two pathways are largely mutually exclusive, as only 20 genomes (15 from Actinomycetes) showed evidence of both pathways, which represents only 0.15% of the 13,168 genomes annotated for MK production. This trend had been previously observed [27], but it was notable that it could be confirmed across such a wide range of bacterial species. Interestingly, we found that the two pathways were intermingled in 47 orders (11.5%) of the 407 orders that produce MK. For further investigation, we selected two of the most populated bacterial lineages that showed several orders with both the Men and futalosine pathways: the *Actinomycetota* (Fig 2B) and the super-lineage previously known as Deltaproteobacteria (Fig 2C). Species trees were reconstructed for both lineages, allowing to map the distribution of the two MK pathways and to analyze their evolutionary dynamics. We observed multiple switches between the two pathways for both lineages (Fig 2B-C), although some lineages preferentially harbored one pathway (*e.g.* Men for *Acidimicrobiia* or futalosine for *Desulfobacterota* Fig 2B-C). This pattern of switching between pathways strongly suggests regular acquisition of one MK pathway by lateral gene transfer alongside the loss of the ancestral pathway, as observed e.g. in *Desulfobacterota* (Fig 2C). However, under certain circumstances, both pathways are maintained, as exemplified by a lineage of Actinomycetes (Fig 2B). Aside from these cases, we also found multiple genomes in which neither pathway was present (S2 Fig and S3 Fig). Overall, the evolutionary history of the two MK pathways is surprisingly highly dynamic, which had not been previously documented. The reasons for frequent exchanges of MK pathways or for the maintenance of both pathways in specific lineages remain to be investigated and may reflect specific metabolic adaptations.

**Fig 2.**
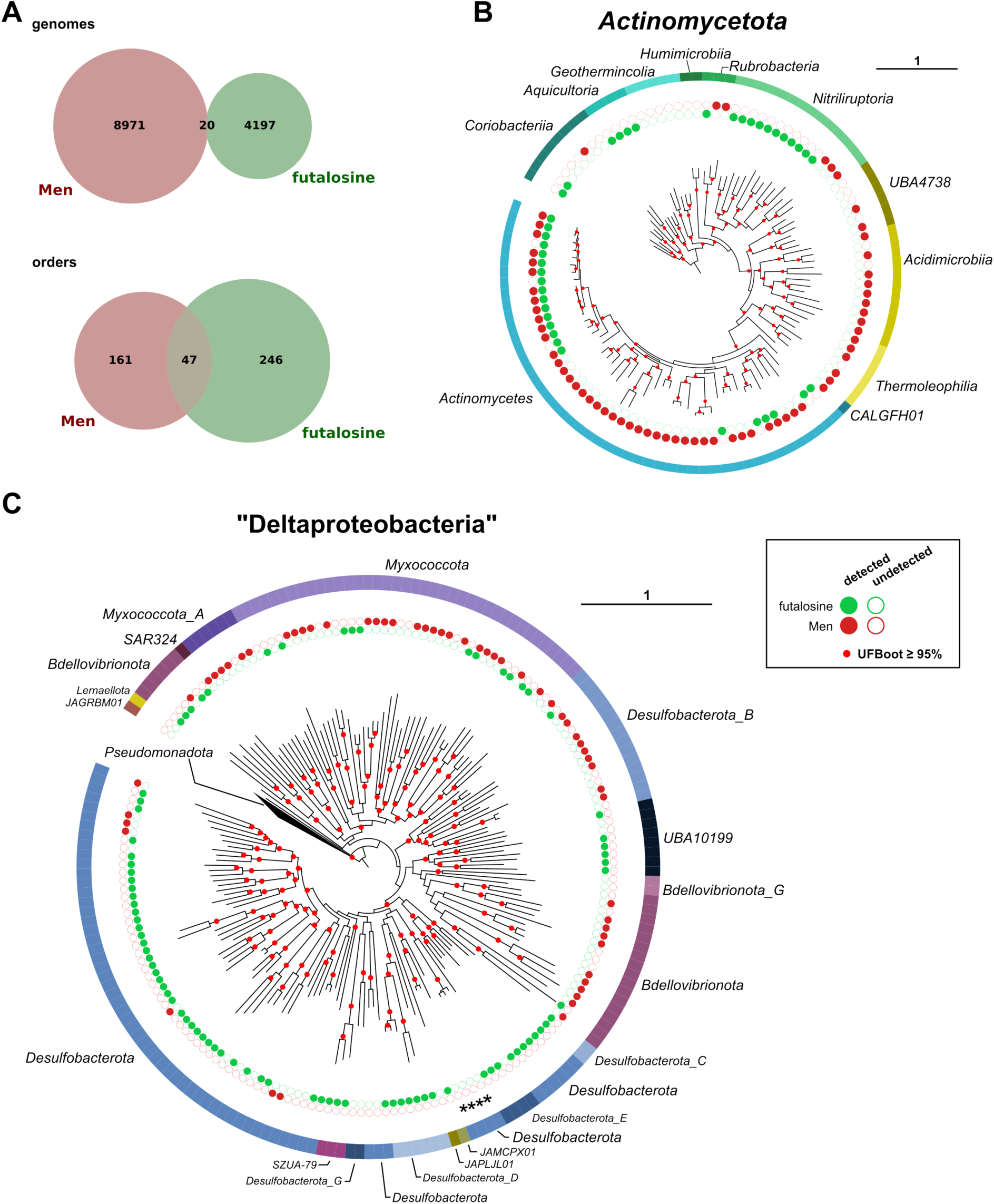
Observation of the disjoint presence of Men and futalosine pathways. **(A)** Genome-level and order-level counts for the presence of the Men and futalosine pathways. **(B-C)** Maximum-likelihood species trees of *Actinomycetota* (B) and phyla corresponding to the former Deltaproteobacteria (C). The presence of the Men and futalosine pathways is annotated on the trees, along with the taxonomy at the class level for *Actinomycetota* and at the phylum level for the “Deltaproteobacteria”. Stars indicate the prediction of UQ in the class GWC2-55-46. The trees were constructed from the concatenation of 120 GTDB markers of 108 *Actinomycetota* and 185 “Deltaproteobacteria” genomes, respectively.

UFBoot values over 95% are depicted by red dots and the tree scale is expressed in substitutions per site.

### Discovery of an ancient UQ pathway in a lineage outside of Pseudomonadota

UQ is largely dominant in *Pseudomonadota* and has only been reported in one instance in another bacterial phylum [49], a finding that is not supported by our genomic analyses (see above). Thus, in bacteria, UQ is currently considered to occur exclusively in *Pseudomonadota* [24]. The class GWC2-55-46 from the *Desulfobacterota* phylum, in which we detected the UQ pathway (Fig 1), was therefore of particular interest. This class is represented in our dataset by nine high quality metagenome-assembled genomes (MAGs) originating from nine distinct environments, including hydrothermal vents, groundwater, sediment and sludge. Contrary to the rest of the *Desulfobacterota*, which is mostly composed of MK producers (Fig 2C and S3 Fig, S1 Table), none of these MAGs showed any sign of a MK pathway. Instead, we were able to annotate a complete UQ pathway comprising *ubiA, -B, -C, -D, -E, -G, -K, -X*, and the three genes involved exclusively in O_2_-independent UQ biosynthesis: *ubiT, -U, and -V* (Fig 3A, S1 Table). However, in genome GCA_024255045.1, *ubiG* was absent and *ubiD* was located at the boundary of a contig. Although the genomes were not fully assembled and were therefore not contiguous, the conserved genomic organization of the UQ genes was evident across the class and was the most compact ever observed [24], with two well-conserved loci comprising the *ubiCADG* and *ubiVKBET* genes (Fig 3A). The closest known organization was that of *Magnetococcia* within *Pseudomonadota*, with several shared colocalized genes, most of them being more scattered in the genome of *E. coli* (Fig 3A). Interestingly, one of the genomes (GCA_001595385.3) harbored a *rquA* gene that suggested RQ production from UQ. To test the functionality of the UQ pathway discovered in *Desulfobacterota*, all the gene candidates from the GCA_001595385.3 genome were expressed in relevant *E. coli ubi* genes mutant strains. Except for *ubiK*, the expression of all the tested heterologous genes resulted in an increased production of UQ (Fig 3B-D). Therefore, we conclude that these genes have the expected function and collectively constitute a complete UQ biosynthesis pathway in the GCA_001595385.3 genome (Fig 3E). The functionality of the *rquA* gene remains questionable, as RQ production was not observed in the wild-type *E. coli* strain expressing *rquA* (data not shown). Overall, the data strongly support the capacity of *Desulfobacterota* from the GWC2-55-46 class to synthesize UQ via an O_2_-independent pathway. This is the first evidence of UQ biosynthesis in bacteria outside the *Pseudomonadota* phylum.

**Fig 3.**
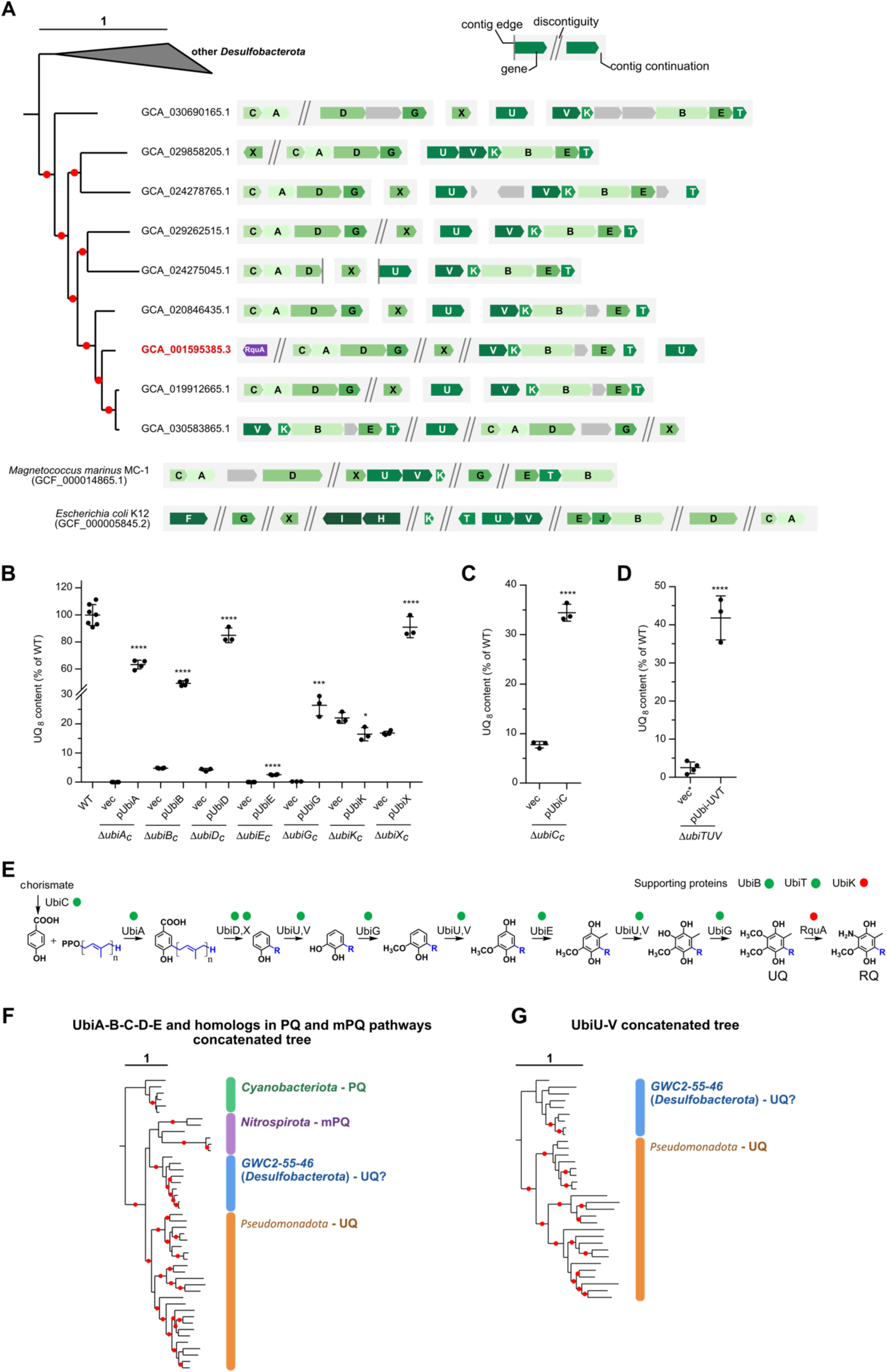
Characterization of a UQ pathway in the genomes of class GWC2-55-46. **(A)** Species tree of GWC2-55-46 and genomic organization of genes involved in UQ biosynthesis. The tree was constructed as the one in Fig 1, with all the high-quality genomes from the GWC2-55-46 class and five close *Desulfobacterota* members selected as outgroups. The UQ/RQ pathway genes were visualized using the GenomeViz Python library (v0.4.4, https://moshi4.github.io/pyGenomeViz) in a custom script, both along the tree and for two species of *Pseudomonadota*, *Magnetococcus marinus* and *E. coli*, which display different genetic organizations. The genome which genes were selected for heterologous expression in *E. coli* is highlighted in red. **(B-D)** Quantification of UQ8 content in *E. coli* Δ*ubi* mutant strains expressing candidate Ubi proteins from GCA_001595385.3 or containing an empty pBAD plasmid (vec), or an empty pTU2b-RFP plasmid (vec*). Quantifications are expressed as percentages of the control values, which correspond to the UQ8 content of the wild-type strain. Mean ± SD (n = 3-5). **** p < 0.0001 ; *** p < 0.005 ; * p < 0.03 (by unpaired Student’s t-test comparing cells containing the plasmid-encoding gene to cells containing the empty plasmid). **(B-C)** Cells were grown aerobically overnight at 37°C in LB medium (A) or M9 minimal medium supplemented with 0.4% glucose (B). **(D)** The Δ*ubiUVT* strain was grown in LB medium under anaerobic conditions. **(E)** Results of the heterologous expression assay of UQ/RQ O2-independent pathway genes from GCA_001595385.3. A green dot next to the protein indicates successful complementation in *E. coli*, while red dot symbolizes a negative result. **(F)** Concatenated trees of UbiA-B-C-D-E and their homologs in the PQ and methyl-PQ pathways. **(G)** Concatenated trees of UbiU-V. All trees were inferred using IQ-TREE [50]. Individual trees are displayed in S4 Fig and S5 Fig. UFBoot values over 95% are depicted by red dots and the tree scale is expressed in substitutions per site.

To investigate the origins of the *Desulfobacterota* UQ pathway, we constructed phylogenetic trees with homologs from various evolutionarily related quinone pathways. We included proteins involved in the synthesis of PQ in *Cyanobacteriia*, UQ in *Pseudomonadota,* and the recently discovered methyl-PQ in *Nitrospirota* [25]. The obtained trees robustly delineated the proteins involved in producing each of the three quinones (Figs 3F and S4 Fig). The UQ proteins from GWC2-55-46 (*Desulfobacterota*) formed a highly supported independent clade that branched as a sister group to the UQ proteins from *Pseudomonadota,* rather than branching among the *Pseudomonadota*. A similar topology was observed for the UbiU-V proteins (Fig 3G and S5 Fig). The precise position of mPQ proteins relative to the two UQ clades could not be determined with certainty, as the tree was not fully resolved. However, the mPQ proteins fell within a well-supported monophyletic group together with UQ proteins from GWC2-55-46, forming a sister-lineage to *Pseudomonadota* UQ proteins (Fig 3F), as previously observed [25]. Taken together, these data rule out the possibility of the GWC2-55-46 class acquiring the UQ pathway from one of the known lineages of *Pseudomonadota*. However, they do not entirely exclude the possibility of lateral gene transfer from an ancient *Pseudomonadota* lineage. Since the *Pseudomonadota* probably innovated UQ over two billion years ago prior to diversifying [24], we propose that the UQ pathways of the GWC2-55-46 class and extant *Pseudomonadota* diverged early on.

### A diversity of quinone molecular features revealed from text-mining data

Although the genomic annotation of biosynthetic pathways and the extraction of quinone data from the literature showed very consistent patterns (Fig 1), these two approaches are fully complementary, as some “rare quinones” cannot be predicted from genomes and are only accessible through experimental data. Mapping the results of the text-mining along the bacterial tree of life revealed that *Pseudomonadota* have the greatest diversity of quinones, including MK, DMK, (D)MMK, UQ, RQ, and methylene-UQ, a variant of UQ with a modified chain (Fig 4). *Aquificae* were shown to produce methionaquinone, which is otherwise found in some archaeal lineages [14]. A few lineages were shown to produce the overmethylated (di)methylmenaquinone (*Thermincolia*, *Coriobacteriia*, a few *Gammaproteobacteria*), in agreement with previous results [29,51] (S2 Table).

**Fig 4.**
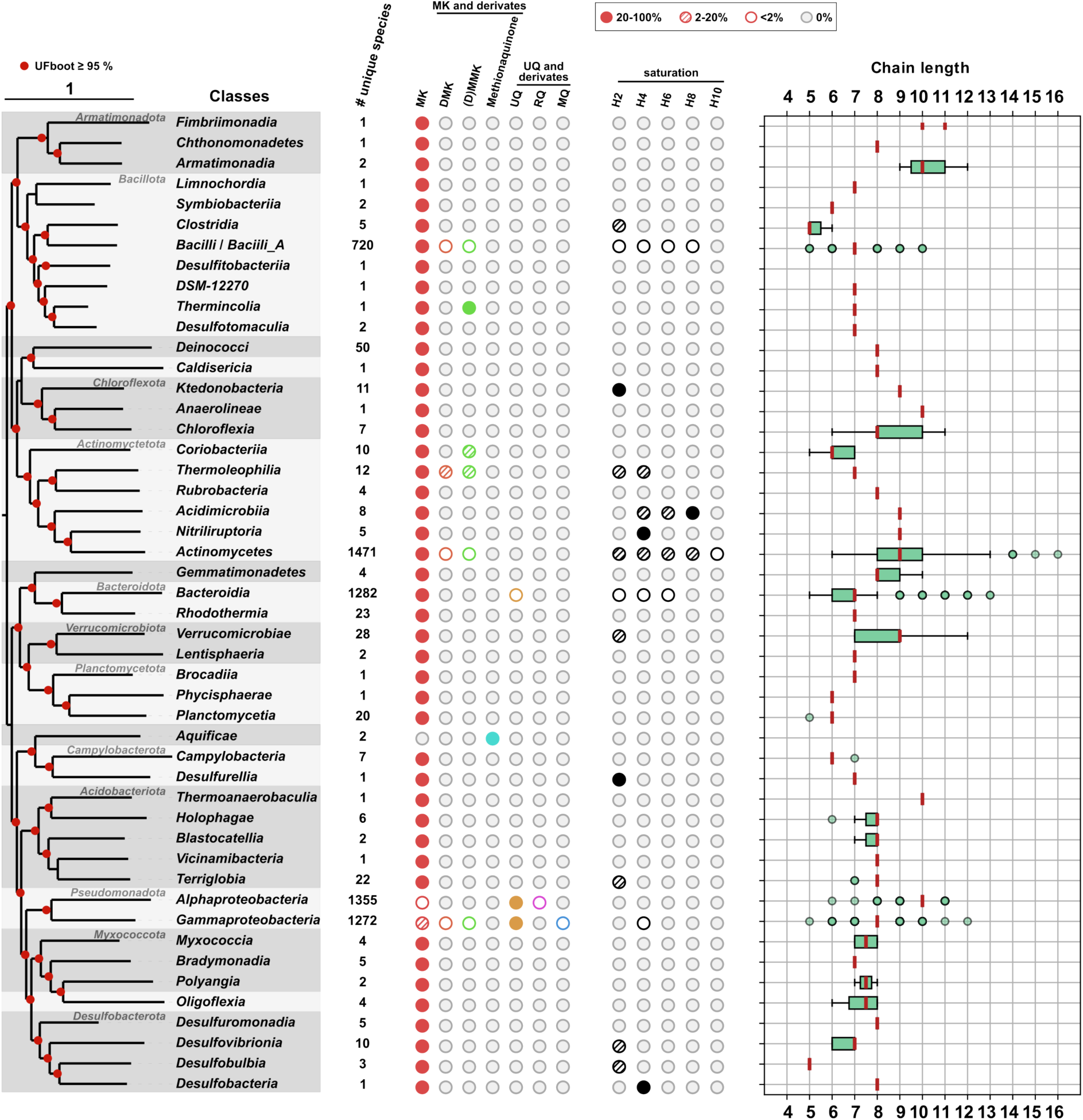
Distribution of quinones chain length and saturation along a phylogenetic tree of *Bacteria*. The maximum-likelihood species tree was constructed using the concatenation of 120 GTDB markers and one genome from each of the 48 classes present in the text-mining dataset. It was rooted as in Fig 1. Phyla are differentiated by shades of gray, those with multiple classes have their names indicated. The following data from the text-mining dataset is displayed along the tree from left to right for each class: the number of unique species, the distribution of each quinone, the distribution of the number of saturations on the isoprenoid chains, and the distribution of chain lengths in the form of boxplots. The boxplots represent the median (Q2) in red, with 50% of the data covered by the box (Q1-Q3), and the ends of the whiskers corresponding to the minimum and maximum chain lengths within 1.5 times the interquartile range before Q1 and after Q3. Outliers are represented by small circles. The edge of the circle is thick when two or more outliers share the same value.

Another piece of information that cannot be obtained from genomic annotation, is the length and saturation of the isoprenyl chains of quinones. This information was extracted from the literature data: chain size was specified in 8,950 cases, and saturation in 1,758 cases (S2 Table and S2 Text). The chain length varied greatly across *Bacteria*, with most quinones displaying chains composed of 5 to 11 isoprenyl units (Fig 4). Some well-covered classes, such as *Deinococci* (50 species) and *Bacilli* (719 species) displayed rather conserved chain sizes of 8 and 7, respectively. Within *Pseudomonadota*, quinones from the *Alphaproteobacteria* class (1,355 species) had mostly a chain length of 10 isoprenyl units, while *Gammaproteobacteria* (1,272 species) had mostly 8 (Fig 4). This strong conservation at the class level contrasted with the situation observed for example in *Actinomycetes* (1,471 species), which displayed a large diversity of chain sizes ranging from 6 to 16 (median size: 9). This variability was observed down to the family level with the *Microbacteriaceae* displaying an unusual diversity of chain lengths (S6 Fig). Within individual species, quinones were commonly reported with side chains varying by up to two isoprenyl units (S2 Table), likely because of the plasticity of the polyprenyl diphosphate synthases that produce chain precursors [12]. *Microbacterium algeriense* was an unusual case, as it was reported to contain MK_9_-MK_13_ [52].

Partial saturation of the side chain of quinones (reduction of some C-C double bonds to single bonds) was observed almost exclusively for MK (1,199 cases) and DMK (12 cases), with degrees of saturation ranging from one (H_2_) to five (H_10_). The 1,211 cases corresponded to 1,161 species distributed in 15 classes (Fig 4, S2 Table). Monosaturated MKs were common in *Ktedonobacteria* (MK_9_-H_2_ in 10/11 species) and *Desulfovibrionia* (MK_6_-H_2_ or MK_7_-H_2_ in 5/10 species), whereas disaturated MKs were prevalent in *Thermoleophilia* (MK_7_-H_4_ in 8/12 species) and *Nitriliruptoria* (MK9-H_4_ in 5/5 species). MKs from *Acidimicrobiia* showed an even higher degree of saturation, with two species containing MK_9_-H_6_, and 7 species containing MK_9_-H_8_. Partially saturated (D)MKs were the exception in *Bacilli* (found in 7/719 species) but the rule in *Actinomycetes* (1,106 species with partially saturated (D)MKs and 365 with unsaturated (D)MKs). Some species, such as *Actinokineospora spheciospongiae* and *Actinomadura parmotrematis* showed an extreme variability in the degree of saturation with MK_9_ ranging from H_0_ to H_8_ [53,54]. In summary, our text-mined quinone dataset is the most comprehensive compilation to date, offering an unparalleled view of the diversity of quinone molecules produced by *Bacteria*.

## Discussion

The present study provides a comprehensive overview of quinone distribution in *Bacteria* by combining the genomic annotation of known quinone pathways with experimental quinone characterizations from the literature. The two types of data were highly consistent, with approximately 98% of quinones produced matching the genomic predictions. While a few discordant cases may suggest potential methodological errors (see S1 Text), others highlight knowledge gaps that require further investigation. For instance, *Gloeobacter violaceus* and *Anthocerotibacter panamensis*, which belong to a deep-branching lineage of *Cyanobacteriia*, are known to produce MK or phylloquinone, yet show no genomic evidence of a biosynthetic pathway [55,56]. This finding suggests the existence of an alternative MK pathway. It is evident that the prevailing function of quinones in bioenergetics and other cellular processes [1,2] renders the unified quinone dataset presented here instrumental in enhancing our comprehension of bacterial diversification. While some lineages exhibited a homogeneous content of quinone types, consistent with the prior use as chemotaxonomic markers, others displayed surprising diversity in terms of chain length and saturation (Fig 4). The large-scale dataset provides an opportunity to investigate the functional relevance of different quinone types by probing the correlations between quinone composition and traits such as membrane lipid composition, the genomic content of bioenergetic enzymes, and ecological niches. Furthermore, the dataset enables the identification of genomic determinants responsible for quinone molecular diversity. For instance, comparative genomic analyses could identify the prenyl reductases involved in quinone tail saturation, and artificial intelligence models could be trained to improve prediction of the chain length produced by polyprenyl diphosphate synthases [12].

Genomic annotation revealed a UQ pathway in the GWC2-55-46 class of the phylum *Desulfobacterota*. This finding was unexpected, given that UQ had previously been identified only once outside the phylum *Pseudomonadota*, in a species of predatory bacteria from the phylum *Bdellovibrionota* [49]. However, in this case, genomic analyses could not conclusively demonstrate the presence of a UQ biosynthetic pathway, as only three homologs were identified [49]. Conversely, all nine MAGs from the GWC2-55-46 class contained 10–11 putative UQ biosynthesis genes, all of which were experimentally validated in *E. coli* mutants, except for *ubiK* (Fig 3B-E). This *Desulfobacterota* UQ pathway provides further evidence regarding the nature (O_2_-dependent or O_2_-independent) of the ancestral UQ pathway. Indeed, GWC2-55-46 genomes exclusively contain the O_2_-independent UQ pathway, with the common genes (*ubiKBE*) intermingling at the same loci as the genes of the O_2_-independent pathway (*ubiTUV*) (Fig 3), as only observed thus far in *Magnetococcia* [23]. The phylogenetic placement of the *Desulfobacterota* pathway outside the currently characterized *Pseudomonadota*, in conjunction with the basal position of the *Magnetococcia* pathway, lends support to UQ biosynthesis initially operating via the O_2_-independent pathway. This hypothesis is consistent with the biogeochemical context proposed for the emergence of UQ, either around the Great Oxidation Event [11,28] or even prior to it [25].

Another notable finding was the high rate of exchange between the two MK pathways within several phyla (Fig 2). Since both pathways produce the same molecule from the same precursor, what evolutionary advantage is there for distinct bacteria to use either the futalosine or Men pathway? Factors such as differences in regulation, cofactor requirements, or adaptation to ecological niches may be involved. Following the establishment of reliable annotation methods for both pathways across bacterial genomes [24], the investigation of such hypotheses is now possible. The utilization of reliable annotation tools will also facilitate the investigation of which MK pathway evolved first. Based on phylogenetic analyses of a smaller genomic dataset, Zhi and colleagues proposed that the futalosine pathway is the oldest [27]. This hypothesis is supported by the much broader taxonomic distribution of the futalosine pathway compared to the Men pathway (Fig 2A). However, definitive answers regarding the evolution of the MK pathways require further investigation, as our study highlighted their intricate distribution across several bacterial lineages (S2 Fig) and their frequent loss and lateral transfer. These evolutionary events reflect a highly dynamic repertoire of quinone pathways, whose physiological relevance across the bacterial kingdom is yet to be determined. Recent research conducted on the *Pseudomonadota* has indicated that the Men pathway and the O_2_-independent UQ pathway are associated with adaptations to microoxic and anoxic environments [24]. This finding suggests that large-scale investigations of quinone repertoire variations may help unravel metabolic adaptations. Phyla containing species with either low-potential (MK) or high-potential (UQ, mPQ) quinones are expected to provide particularly valuable insights, given the significant implications of such quinone differences for bioenergetics. It has recently been demonstrated that the *Nitrospirota* display this specificity, with early-branching lineages consisting of anaerobes that produce MK, and more recent lineages composed of aerobes that produce mPQ [25]. The innovation of this high potential quinone may provide an advantage in oxic environments populated by *Nitrospira* species and could be involved in the respiratory chain and reverse electron transport [26]. The *Desulfobacterota* phylum, which comprises numerous anaerobic sulfate-reducing species that produce MK, constitutes a novel instance of acquisition of a high potential quinone, as species in the GWC2-55-46 class contain exclusively the O_2_-independent UQ pathway. Interestingly, members from this class have been proposed to harbor high affinity O_2_ oxidases, which suggests that these bacteria are using O_2_ as an electron acceptor in micro-oxic conditions [57,58]. The future isolation or culture enrichment of members of this class could validate UQ production and provide further insight into their metabolism.

The present study emphasizes the importance of investigating the dynamics of the quinone repertoire across bacteria and of correlating it with the diversification of energetic metabolisms and ecological adaptations. The quinone compendium provided here establishes the foundation for a comprehensive understanding of the role of quinone diversification in the evolution of bacterial metabolism. It is anticipated that the dataset will be augmented by ongoing experimental characterization of quinones produced by newly identified species.

## Material and Methods

### Genomic dataset and annotation of quinone pathways

We downloaded 26,264 protein files corresponding to the high-quality reference genomes from release 220 (April 2024) of the Genome Taxonomy Database (GTDB) along with the associated taxonomy [42,59,60]. We used Hidden Markov Model (HMM) profiles published in previous studies to identify the genetic potential for UQ, RQ, MK, and PQ biosynthesis [24,61]. The method to infer the presence of the UQ, RQ, and MK pathways was the same as in a previous study [24]. Overall, we inferred the presence of these pathways in the genomes when at least the minimum number of proteins required for the pathway could be annotated (3, 5, 7, 6, and 6 proteins respectively for the PQ, UQ O_2_-dependent, UQ O_2_-independent, Men, and futalosine pathways, see S1 Table). For the mPQ pathway, as profiles are not yet available for this elusive biosynthetic pathway, the distribution’s results were based on our recent investigation of the gene set we showed to be involved in mPQ biosynthesis [25].

### GTDB Taxonomic assignments

To compare bioinformatics quinone annotations with text-mining extractions, the species identified in the text-mining approach were reassigned to the GTDB taxonomy via their NCBI taxid. As a single NCBI taxid could correspond to multiple taxa in the GTDB, the most relevant taxonomy was selected based on the species name, prioritizing simpler and more closely related names. The following priority order was used: (i) an exact match for the presented species name, (ii) a slight variation in the genus name (e.g., ‘*Rhodococcus_D*’ instead of ‘*Rhodococcus*’), (iii) a match for the genus but slight variation in the specific name of the species binomial nomenclature (with and without variants such as ‘_A’, ‘_B’, ‘_C’). Whenever multiple taxonomies were still possible, (iv) the simplest and first option by alphabetical order was chosen (e.g., *‘Flavobacterium oncorhynchi_A*’ over ‘*Flavobacterium oncorhynchi_B*’). If multiple taxonomies still remained, (v) a random selection was made. Ultimately, the NCBI taxid of 6,459 species could be matched to the GTDB taxonomy, while 2,368 NCBI taxid could not be matched (S2 Table).

### Species trees

To build species trees, alignments provided by the GTDB (release 220) for its 120 phylogenetic markers were downloaded and concatenated for the genomes of interest (see below for the description of the different genomic samples). IQ-TREE (v 2.2.0.3) was used to build the concatenated alignment and to infer a maximum likelihood tree [50] using the options ’-p’ and ’-B 1000’ to calculate branch supports with ultrafast bootstraps (UFBoot) [62]. ModelFinder was employed to determine the best model for each marker individually [63].

Several samplings were performed for the reconstruction of phylogenetic species trees: (i) A total of 751 bacterial genomes were selected for the tree presented in Fig 1 and S2 Fig. To illustrate quinone diversity, we randomly selected one genome per class when all genomes within the class had an identical quinone pathway content. Otherwise, we randomly selected one genome per order. Genomes from species identified during the text-mining step as containing rare types of quinones were preferentially selected over random choices. (ii) One genome from each class containing at least one species with a described quinone was selected for the tree presented in Fig 4. (iii) To construct the *Actinomycetota* tree (Fig 2), we sampled between one and four genomes per order, prioritizing genomes that displayed diversity in their quinone biosynthesis pathways. Genomes containing both the Men and futalosine pathways were oversampled. (iv) We applied the same sampling approach to construct the tree for genomes of organisms previously gathered in the Deltaproteobacteria and Bdellovibrionia phyla (Fig 2). These organisms now correspond to a monophyletic group of phyla, which includes mostly *Myxococcota*, *Bdellovibrionota*, and *Desulfobacterota*. Five genomes from the sister group *Pseudomonadota* were used to form an outgroup. (v) The nine high quality genomes from the GWC2-55-46 class, along with five genomes from closely related *Desulfobacterota*, were used to build the tree presented in Fig 3A. The genomes used to construct the various trees are listed in S3 Table.

### Phylogenetic analysis of the quinone biosynthesis proteins

To analyze UQ biosynthesis genes in the GWC2-55-46 class, we constructed concatenated trees for UbiA-UbiB-UbiC-UbiD-UbiE and UbiU-UbiV (Fig 3F-G). We used a subset of protein sequences from the phylogeny constructed in Chobert *et al.* [24], as well as sequences from the nine GWC2-55-46 genomes and homologs in the mPQ (MpqA, MpqB, MpqC, MpqD, MpqE [25]) and PQ (PlqA, PlqC, PlqD [19,25]) pathways. First, individual trees were constructed from a multiple sequence alignment (MSA) obtained with MAFFT using the l-ins-i algorithm (v7.487) [64]. Sites filtering of the MSA was performed using BMGE (v2.00) using the default parameters and the BLOSUM30 matrix [65]. The maximum likelihood trees were generated using IQ-TREE (v2.2.0.3) with the best evolutionary model selected by ModelFinder [50,63]. Branch supports were calculated with 1,000 iterations of UFBoot and SH-aLRT [62,66]. Concatenated trees were built with the ’-p’ option, 1,000 UFBoot iterations, and ModelFinder on the partitioned MSA.

### Text-mining from the abstracts

The annual PubMed baseline database, dated 20 December 2023, was downloaded. A custom script was used to extract the abstracts and related metadata from the following journals: *Antonie van Leeuwenhoek*, *International Journal of Systematic and Evolutionary Microbiology* (formerly known as *International Journal of Systematic Bacteriology*), *Archives of microbiology* and *Systematic and Applied Microbiology.* This resulted in a total of 29,268 articles (S7A Fig). The extracted data for each article included the PubMed ID (PMID), title, abstract, and publication date (year, month). A Python script was developed (available here: https://github.com/TrEE-TIMC/Quinones_pubmed_mining) and used on each article to extract taxonomic information about the studied organism from the title, and information about quinones (quinone type, chain length, and saturation) from the abstract. To extract species names, the text was tokenised and pairs of words were analysed using a sliding window. All pairs of words where the first word started with a capital letter and the second with a lowercase letter were searched as a possible species name using the module NCBItaxa from the Python library ETE3 [67,68]. If the pair of words did not match a species name, NCBItaxa was applied to the first word to identify taxonomic data at any possible level (genus, family, etc.). Finally, NCBItaxa was used to retrieve the complete taxonomy of the species (or genus, family, etc.) identified in the title. Two types of searches using regular expressions were employed to extract quinone-related information: the first search targeted full mentions of quinones in the abstracts (e.g., menaquinone, methylene-ubiquinone), while the second search focused on abbreviated versions of the molecule names (e.g., MK_8_). A filter was applied to remove matches that did not correspond to quinones (often strain names). This filter included removing matches with a chain length greater than 30 units, as well as matches repeated more than four times. Finally, we manually reviewed and corrected, when necessary, cases where: (i) a quinone occurrence was detected but no species was identified, (ii) inconsistencies with the current state-of-the-art were apparent (e.g., very short or very long isoprenyl tail lengths), and (iii) multiple species from different genera were mentioned in the title (S7C-D Fig). As a result, we were able to extract quinone information and their potentially producing species from 9,172 abstracts (S7B Fig).

### Plasmids construction

The nucleotide sequences of the *ubi* gene candidates identified in the GCA_001595385.3 MAG were optimized for expression in *E. coli* and are available in Dataset S1. These genes were synthesized by the companies “GeneCust” and “GenScript” and cloned into the pBAD24i vector downstream of the arabinose-inducible promoter into the NcoI (5’ end) and HindIII (3’ end) restriction sites. The ATG start codon of the candidate *ubi* genes is included in the NcoI site sequence (CCATGG), which requires a G to be the first nucleotide of the second codon. If the second amino acid did not correspond to a codon starting with a G, an alanine codon (GCG) was added before the second codon, as in [24]. The plasmids used in this study are listed in S4 Table.

In the case of the *ubiT*, *ubiU* and *ubiV* gene candidates, the sequences were inserted into the NdeI and BamHI sites of the pBP-ORF plasmid (#72941) from the EcoFlex kit [69]. Care was taken to eliminate BsaI and BsmBI restriction sites in the optimized sequences. Level 1 plasmids containing one transcriptional unit (TU) composed of four DNA bricks (a promoter, a ribosome Binding Site (RBS), the open reading frame and a terminator), were assembled using the NEBridge Golden Gate Assembly Kit (BsaI-HFv2, #E1601, New England BioLabs), in accordance with the manufacturer’s protocol and as described in [23]. The *ubiT* TU was assembled in the plasmid pTU1-A-RFP (#72939), the *ubiU* TU in the plasmid pTU1-B-RFP (#72940) and the *ubiV* TU in the plasmid pTU1-C-RFP (#72941). All assemblies used the J23114 promoter (#72965), the pET-RBS (#72981) and the L3S2P21 terminator (#72999). Positive white clones were selected based on the absence of the red color from the RFP marker, and overnight cultures were used for plasmid DNA extraction (Macherey-Nagel mini-prep kit). The level 2 plasmid containing UbiTUV was constructed by assembling the UbiT, UbiU and UbiV TUs in the pTU2b-RFP vector plasmid (#72959) using the NEBridge Golden Gate Assembly Kit (BsmBI-v2, #E1602, New England Biolabs) in accordance with the manufacturer’s protocol and as described in [23]. The pTU2b-UbiTUV plasmid was verified by ONT Lite Whole Plasmid Sequencing (Eurofins).

### Strains

The strains used in this study are listed in Table S5. The chloramphenicol resistance marker was removed from the Δ*ubiD* strain [70] to yield the Δ*ubiDc* strain, using the plasmid pCP20 as described previously [71]. The Δ*ubiTUV::kan* strain was constructed using one-step inactivation as previously described [72]. In brief, a DNA fragment containing the *kan* gene flanked by the 5’ and 3’ regions bordering the *E. coli ubiTUV* locus was amplified by PCR using pKD4 as a template and the following oligonucleotides: 5wannerTUV: aaagagtagttaaagtt-gttaacaaagtgagctatttacGTGTAGGCTGGAGCTGCTTC; 3wannerTUV: tttcattcccgcctcaacaaaatccgccagttgcagcagcaCATATGAATATCCTCCTTA. Strain BW25113, which carries the pKD46 plasmid, was transformed by electroporation with the amplified fragment, after which Kan^r^ colonies were selected. Replacement of chromosomal *ubiTUV* locus by the *kan* gene was verified by PCR amplification in Kan^r^ clones. The mutation was then introduced into the MG1655 strain via P1 vir transduction [73], followed by selection for Kan^r^ colonies.

### Culture conditions

The empty pBAD24i plasmid and the plasmids containing the gene candidates were transformed into the relevant chemocompetent knock-out strains via heat shock [74]. Transformants were selected on LB plates containing the appropriate antibiotic: ampicillin (50 mg/L), kanamycin (25 mg/L) or chloramphenicol (25 mg/L). 5 mL aerobic cultures were performed at 37 °C in 50 mL glass tubes shaken at 180 rpm. The transformants were cultivated in a medium that contained the appropriate antibiotic and was supplemented with 0.02% (w/v) arabinose to induce expression of the genes cloned into the pBAD plasmids. Lysogeny broth (LB) was used as the growth medium, except for the Δ*ubiC* strain which was cultivated in a synthetic medium (M9) containing 0.4% (w/v) glucose. Autoclaved M9 medium was supplemented with 0.5% (w/v) casamino acids and with 1/100 volume of filter-sterilized solutions of 1mM CaCl_2_, 1mM FeSO_4_, 200 mM MgSO_4_, 1% (w/v) thiamine and 1% (w/v) biotin. Anaerobic cultures of the Δ*ubiTUV* cells were performed in Hungate tubes containing 12 mL of deoxygenated LB medium, as previously described [70]. The cultures were cooled on ice for 30 min prior to centrifugation at 3,200 *g* at 4°C for 10 min. The cell pellets were washed in 1 mL of ice-cold PBS and transferred to pre-weighed 1.5 mL Eppendorf tubes. After centrifugation at 12,000 g at 4°C for 1 min, the supernatant was discarded, and the wet cell weight was determined (5 to 30 mg). The pellets were then stored at -20°C.

### Lipid extractions and quinone analysis

Quinone extraction from cell pellets was performed as previously described [75]. The dried lipid extracts were resuspended in 100 µL of ethanol, and a volume corresponding to 1 mg of cells wet weight was analyzed by HPLC-electrochemical detection-mass spectrometry (ECD-MS) using a BetaBasic-18 column and a mobile phase consisting of 50% methanol, 40%ethanol and 10% of a mixture of 90% isopropanol, 10% ammonium acetate (1 M) and 0.1% (v/v) trifluoroacetic acid. MS detection was performed using an MSQ spectrometer (Thermo Scientific) with electrospray ionization in positive mode (probe temperature 400 °C, cone voltage 80V), as described in [70]. The UQ_8_ peak obtained with electrochemical detection was quantified using a standard curve of UQ_10_ and was corrected for sample loss during extraction based on the recovery of the UQ_10_ internal standard [75].

### Data availability

The data underlying this article are provided as part of the main text and the supplementary data.

## Supporting information

Supplementary Material and Figures (main file)

## Acknowledgments

The authors thank all the teams involved in quinone characterization along the years; they made this work possible. The references used in this study can be found via their PMID in S2 Table. The authors are grateful to the colleagues who reviewed an earlier version of this manuscript and provided valuable feedback: Frauke Baymann, Joël Gaffé, Barbara Schoepp-Cothenet. This project was funded by the French Research Agency (ANR) via the project QUINEVOL (grant agreement ANR-21-CE02-0018).

## Author contributions

SSA and FP designed the research and obtained the fundings. SC performed all the bioinformatic analyses. SK designed a first version of the text mining pipeline under the supervision of SC, NV and SSA. OL, JM and LP performed the experimental analyses and designed the corresponding figures. SC designed all other figures. SC, FP and SSA wrote the original version of this manuscript with contributions from all co-authors. All authors agree with this version of the manuscript.

## Conflicts of interest

The authors declare no conflicts of interest.

## Notes

### Competing Interest Statement

The authors have declared no competing interest.

