## Supplementary Material and Figures (main file) for "Global distribution of isoprenoid quinones across Bacteria"

### Supplementary figures:

S1 Fig. Chemical structure of isoprenoid quinones.
S2 Fig. Phylogenetic tree of bacteria with a detailed view of quinone pathways.
S3 Fig. Phylogenetic tree of « Deltaproteobacteria ».
S4 Fig. Phylogenies of UQ genes: UbiA, UbiB, UbiC, UbiD, UbiE and their homologs in PQ and mPQ biosynthetic pathways.
S5 Fig. Phylogenies of UQ genes: UbiU and UbiV.
S6 Fig. Distribution of chain length variation in Actinomycetes.
S7 Fig. Text-mining data analysis.

### Supplementary texts:

S1 Text. On the cases of discordance between the genomic annotation and the text-mining data.
S2 Text. On the species mentioned multiple times in the text-mining data.

### Supplementary tables:

S1 Table. Genomic annotation of quinone biosynthetic pathways
S2 Table. Text-mining results
S3 Table. Genomes selected for the species trees
S4 Table. Plasmids used in this study
S5 Table. *E. coli* strains used in this study
S6 Table. Cases of discrepancy in species mentioned multiple times

### Supplementary dataset:

S1 Dataset. Optimized sequences of *ubi* gene candidates tested experimentally
S2 Dataset. Phylogenies and supporting data for phylogenies (available on Figshare: 10.6084/m9.figshare.30145930)

### Supplementary figures:

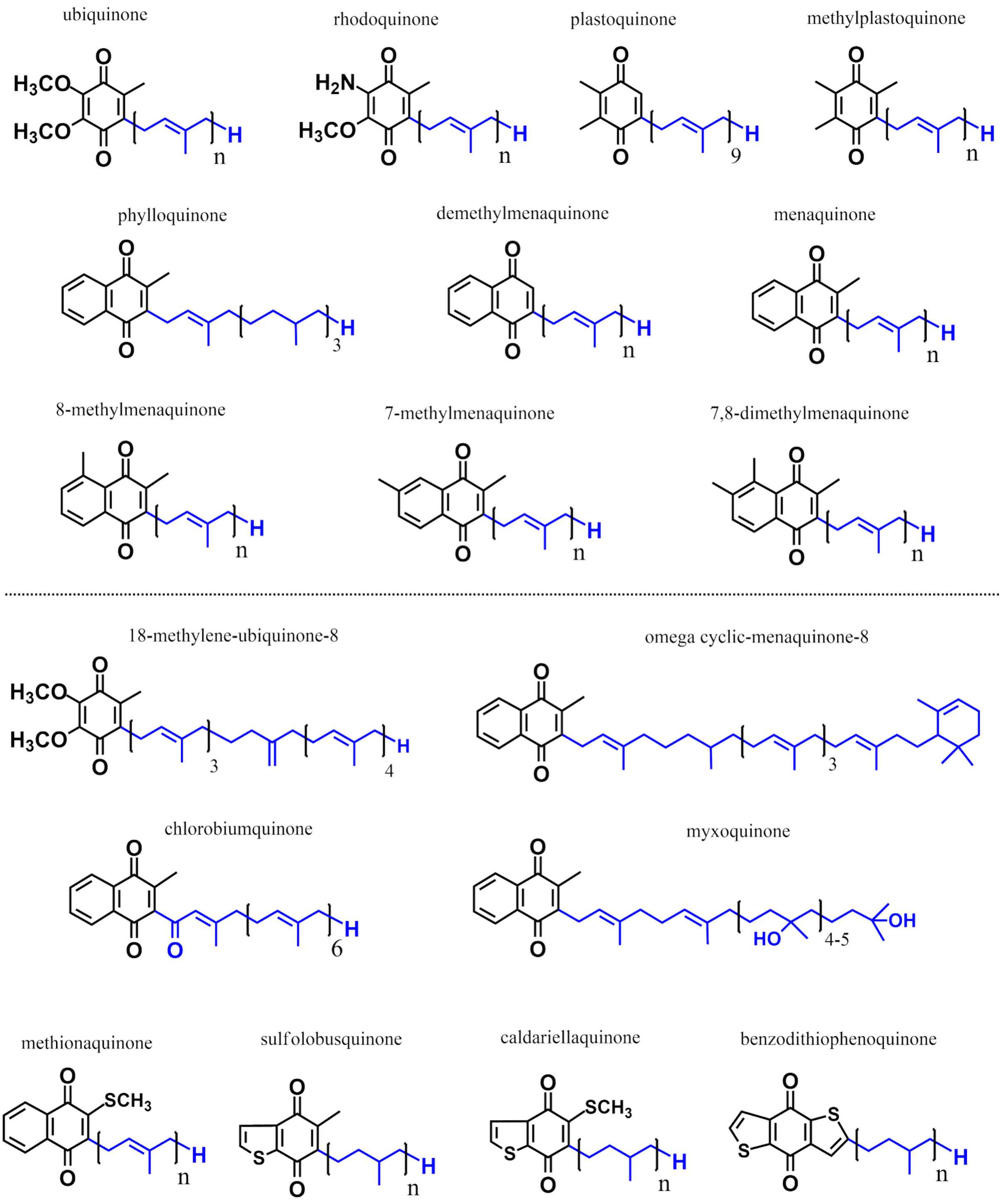


**S1 Fig. Chemical structures of isoprenoid quinones.** The biosynthetic pathways of quinones represented beneath the dashed line remain uncharacterized.

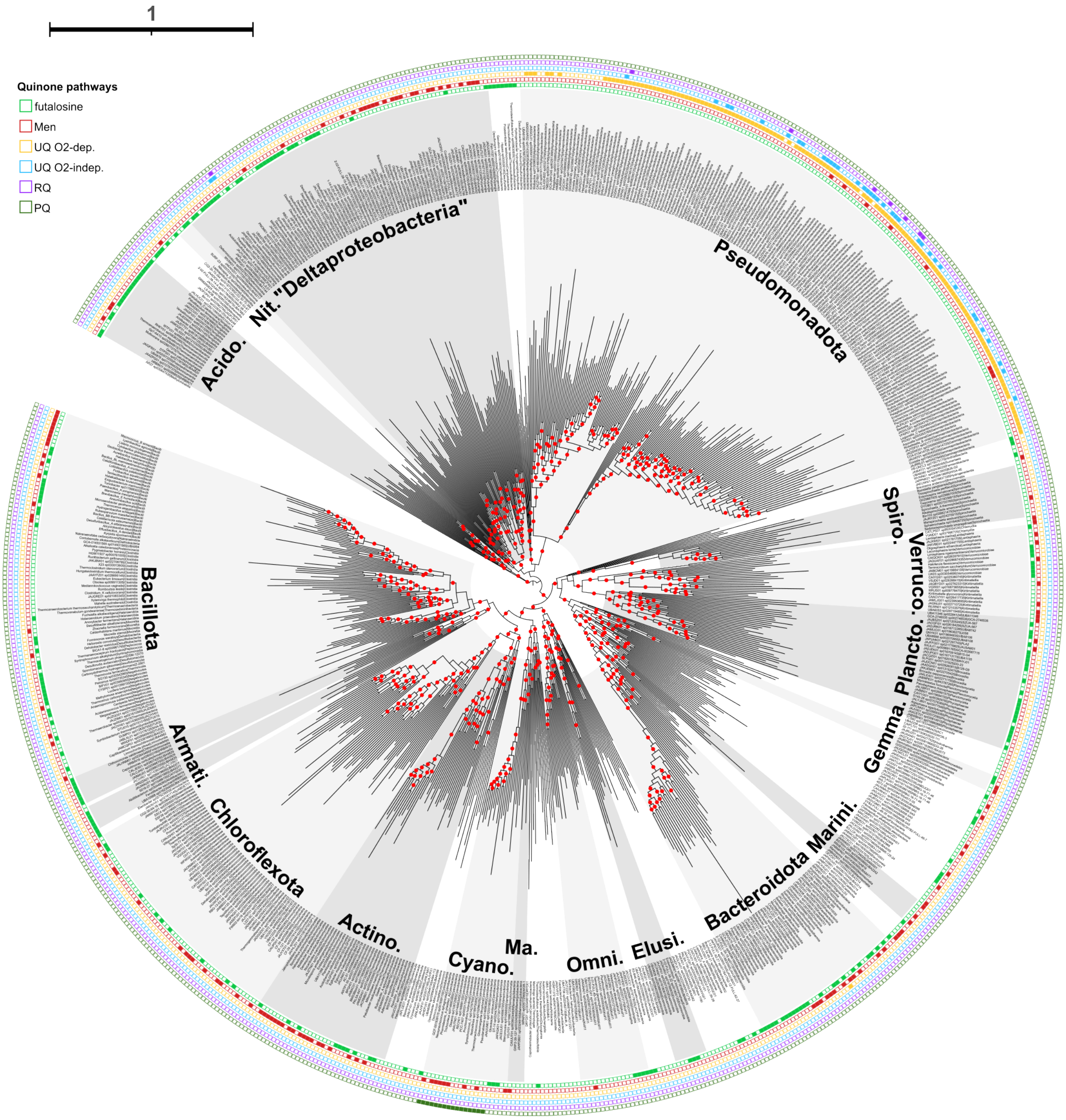


**S2 Fig. Phylogenetic tree of bacteria with a detailed view of quinone pathways.** Tree presented Fig 1. Bootstrap values over 95% are depicted by red circles. The tree was rooted to separate the Gracilicutes from the Terrabacteria. The following selected phyla are labeled on the tree: *Pseudomonadota, Spirochaetota, Verrucomicrobiota, Planctomycetota, Gemmatimonadota, Marinisomatota, Bacteroidota, Elusimicrobiotoba, Omnitrophota, Margulisbacteria, Cyanobacteriota, Actinomycetota, Chloroflexota, Armatimonadota, Bacillota, Acidobacteriota, Nitrospirota* and "Deltaproteobacteria" *(Desulfobacterota, Myxoccocota, Bdellovibrionota* and relatives). The prediction or absence of biosynthetic pathways—MK futalosine (green), MK Men (red), UQ $O_2$-dependent (orange), UQ $O_2$-independent (blue), RQ (purple), and PQ (dark green)—is specified along the tree.

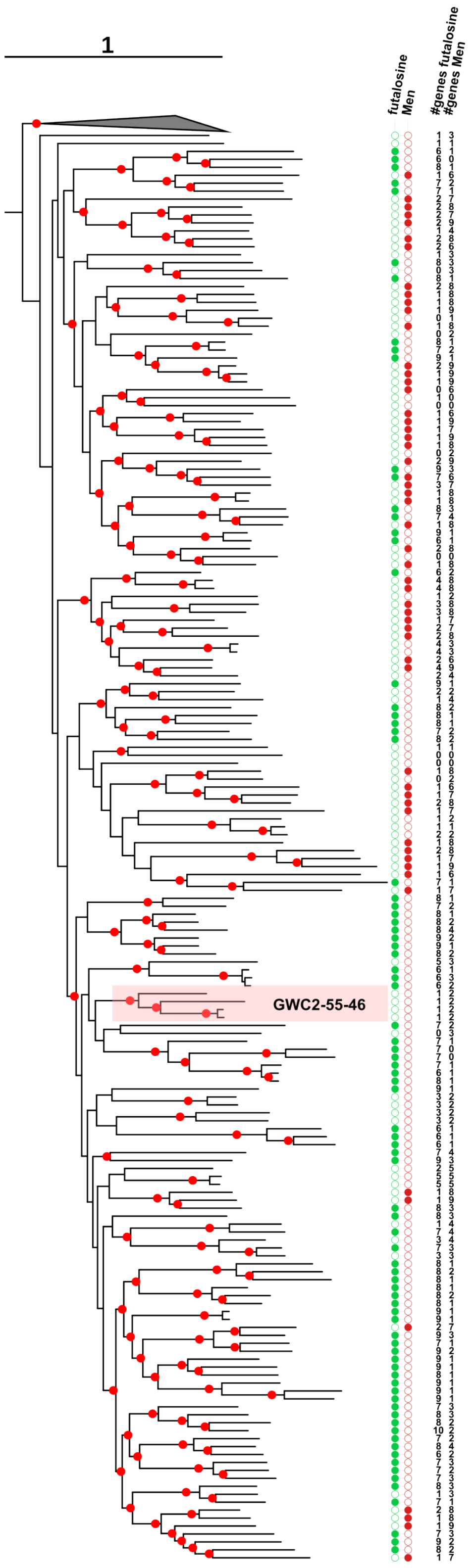


**S3 Fig. Phylogenetic tree of « Deltaproteobacteria ».** Tree presented Fig 2. The prediction or absence of biosynthetic pathways—MK futalosine (green), MK Men (red)—is specified along the tree as well as the number of genes identified for each of the two pathways.

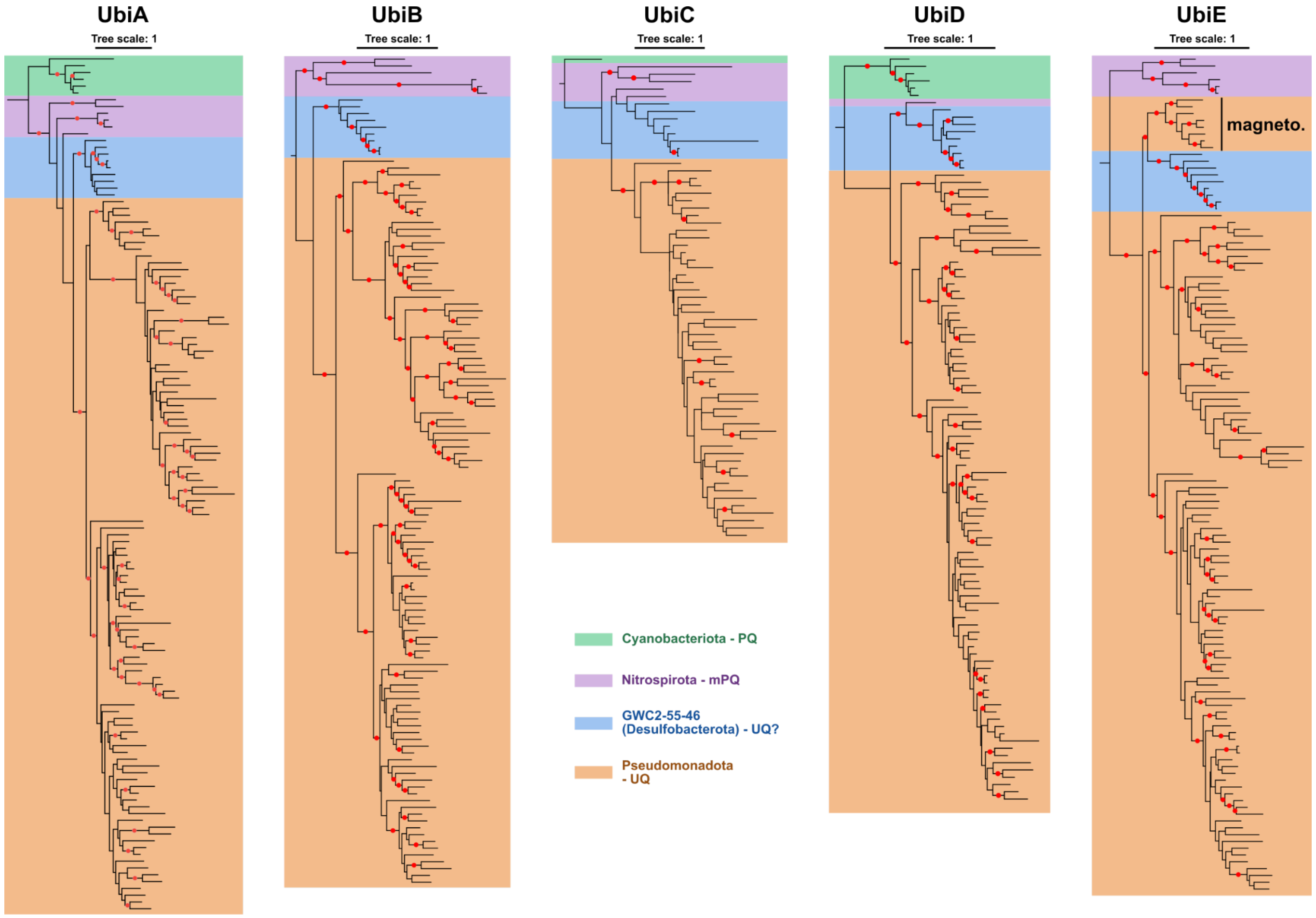


**S4 Fig. Phylogenies of UQ biosynthetic proteins: UbiA, UbiB, UbiC, UbiD, UbiE and their homologs in PQ and mPQ biosynthetic pathways.** The trees of UbiA, UbiB, UbiC, UbiD and UbiE were obtained from the analysis of 247, 422, 79, 474 and 233 aligned positions respectively using IQ-TREE with LG+F+I+I+R7 as the best selected model for UbiA, LG+F+R7 for UbiB, Q.pfam+I+G4 for UbiC, LG+R6 for UbiD, Q.pfam+I+I+R7 for UbiE. The tree scale bar expresses the number of substitutions per site. The branches with very high support (UFBoot ≥ 95%) are indicated by red dots. The trees are rooted using *Cyanobacteria* or *Nitrospirota* sequences as an outgroup when no homologs are found in Cyanobacteria. Overall, tree topologies are close to one another. In the UbiE tree, the group formed by sequences from GWC2-55-46 is positioned as a sister group to a class of *Pseudomonadota*, the *Magnetococcia*.

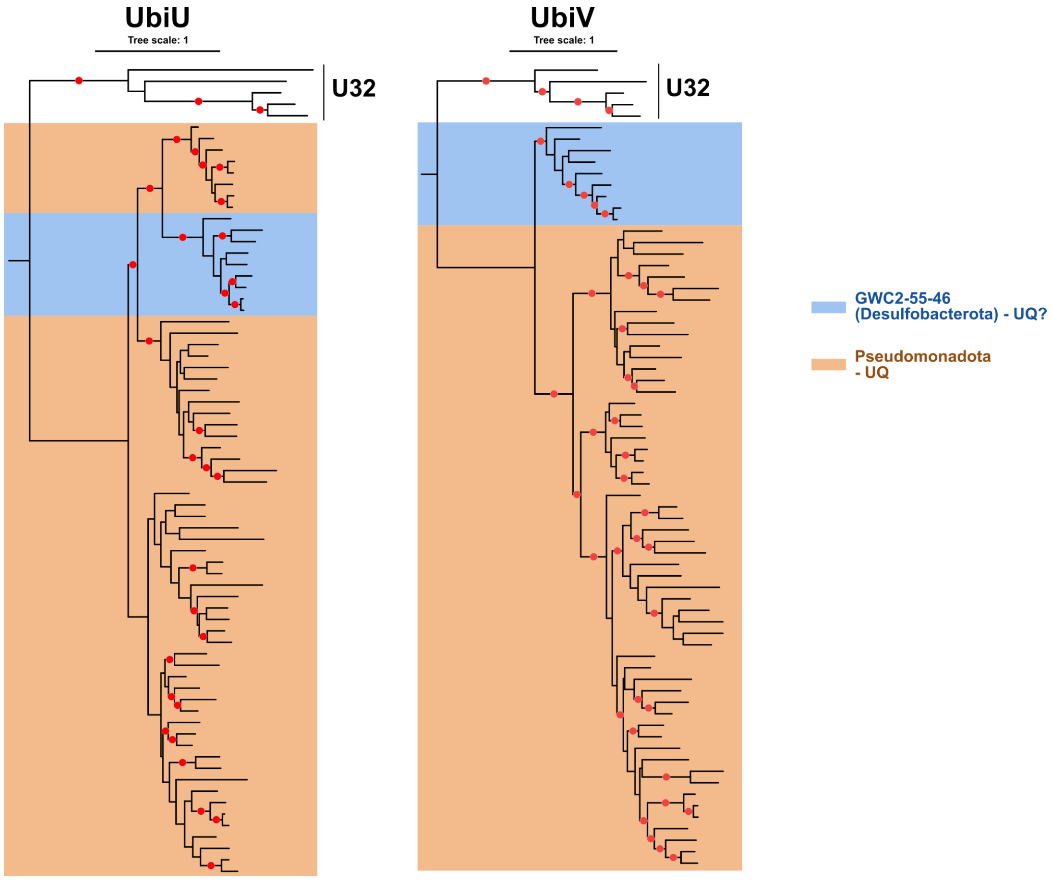


**S5 Fig. Phylogenies of UQ biosynthetic proteins: UbiU and UbiV**. The trees were obtained from the analysis of 302 and 251 aligned positions for UbiU and UbiV using IQ-TREE with Q.pfam+I+G4 as the best selected model for both of the trees. The tree scale bar expresses the number of substitutions per site. The branches with very high support (UFBoot ≥ 95%) are indicated by red dots. U32 protease sequences were used as an outgroup.

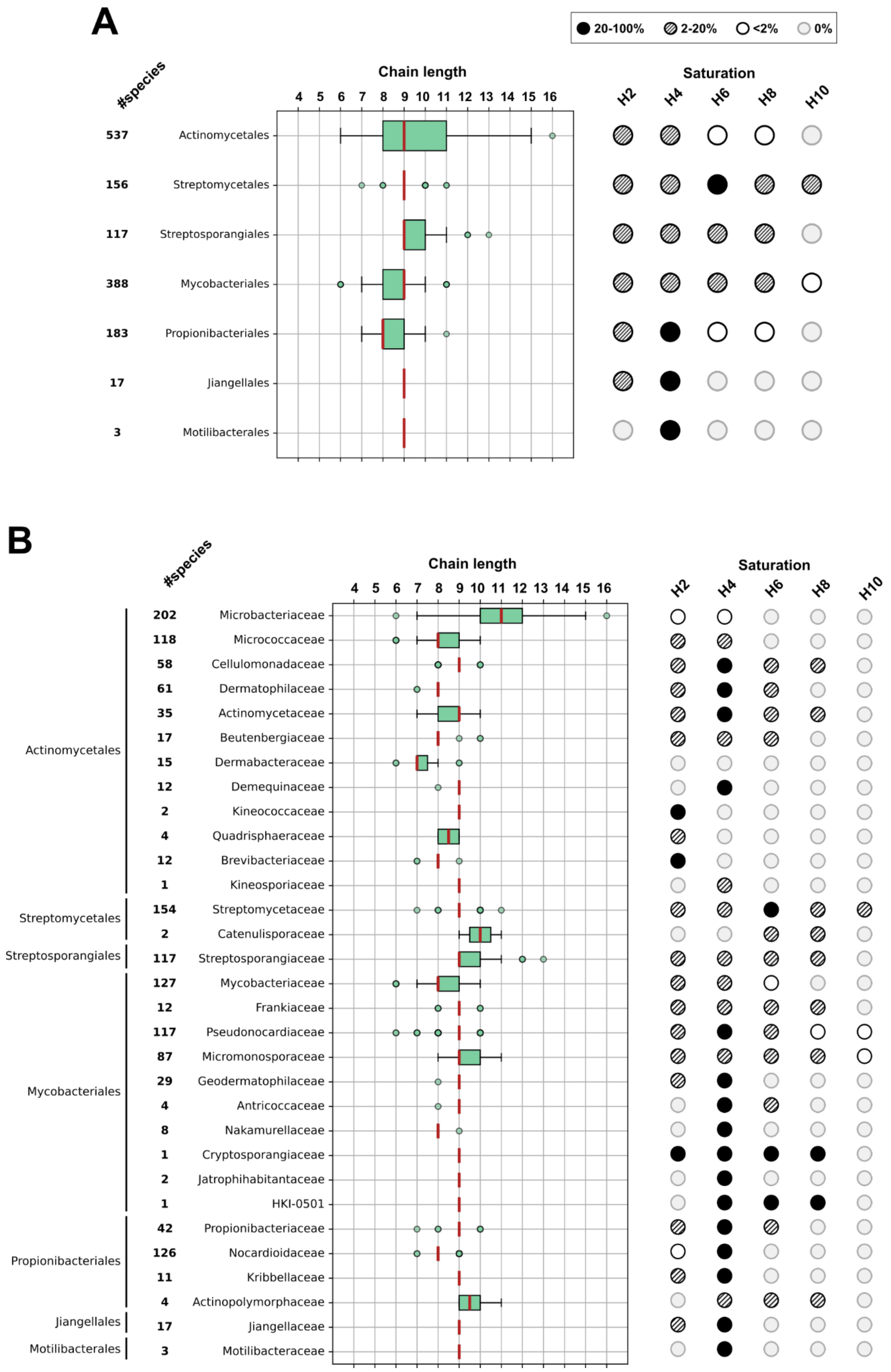


**S6 Fig. Distribution of chain length variation in Actinomycetes.** Chain length variation and saturations at order **(A)** and family level **(B)**. The boxplots follow the typical rule: the median (Q2) is shown in red, 50% of the data is covered by the box (Q1, Q3), and the ends of the whiskers correspond to the minimum and maximum chain lengths within 1.5 times the interquartile range before Q1 and after Q3. Outliers are represented by small circles. Circle edge is thick when two or more outliers share the same value. The number of unique species per order/family is specified along the boxplots.

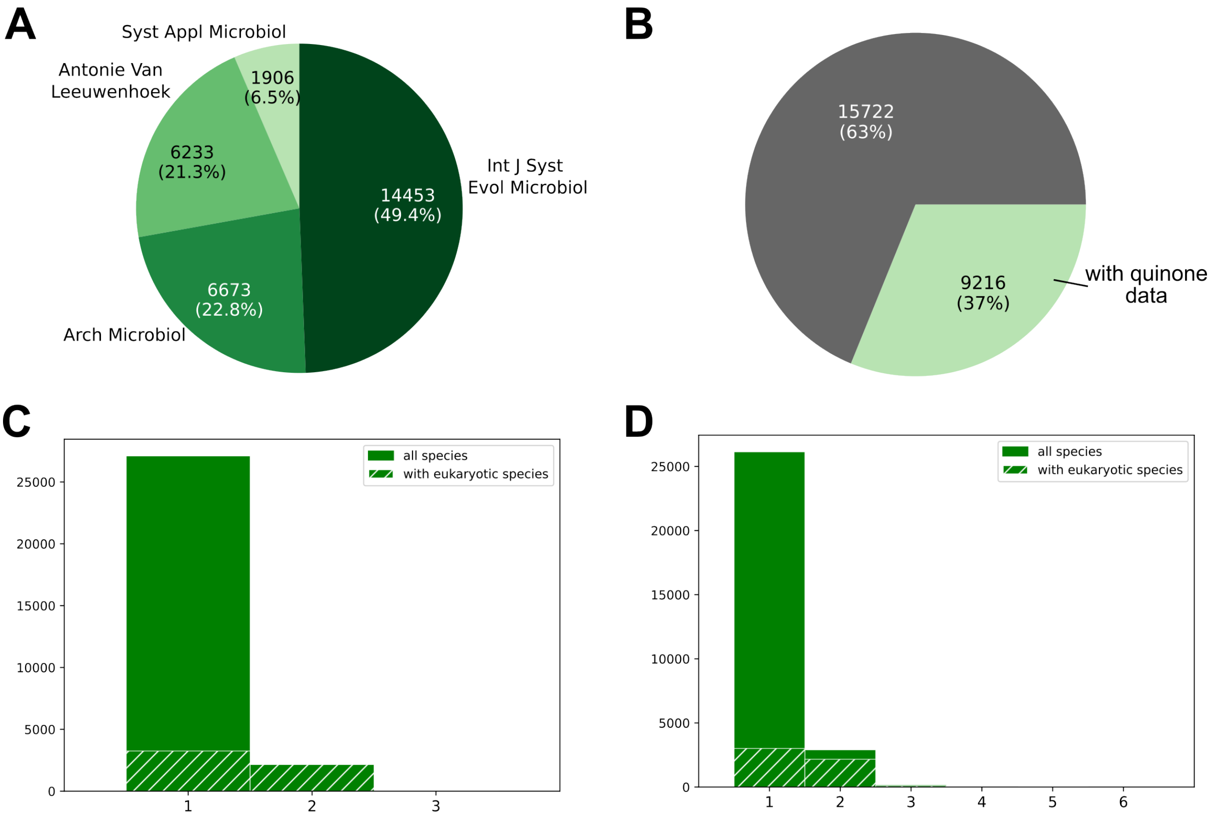


**S7 Fig. Text-mining data analysis**. **(A)** Proportion of abstracts from each journal: Int J System (International journal of systematic and evolutionary microbiology), Arch Microbiol (Archives of microbiology), Antonie Van Leeuwenhoek, Syst Appl Microbiol (International journal of systematic bacteriology) **(B)** Proportion of articles from which information on the type of quinone associated with prokaryotic taxonomic information could be extracted. **(C)** Distribution of the number of different divisions (NCBI taxonomy) per article. The proportion of articles where at least one of the two species is a eukaryote is indicated by hatching. **(D)** Same as C but at genus level. In the context of these articles specialized on prokaryotes, the mention of a eukaryotic species mostly refers to the host species of the bacterium in cases of symbiosis or the source of isolation. In the vast majority of cases, when two genera are mentioned, the other species/genus is a eukaryote. We therefore assume that the description of the quinone refers to the prokaryotic species mentioned.

### Supplementary texts:

### S1 Text:

A few notable cases of discordance between the genomic annotations and the text-mining data were identified (Fig 1, S1 and S2 Tables): MK was measured in *Romboutsia ilealis* (*Bacillota,* PMID: 24480908), *Thermacetogenium phaeum* (*Bacillota_B*, PMID: 10939667), *Thiogranum longum* (*Pseudomonadota,* PMID: 25336721) and *Mesosutterella multiformis* (*Pseudomonadota*, PMID: 30394865), whereas we did not detect any MK pathway in the genomes of these species. In the first three species, no more than two genes involved in MK production were found, suggesting that the genetic potential for MK production is largely incomplete. In contrast, *M. multiformis* harbors five MK-related genes, just below the threshold of six genes required to infer the presence of the MK pathway. In *T. longum,* UQ is annotated, as expected for a *Pseudomonadota*. However, only $MK_8$-$H_4$ and $MK_9$-$H_4$ have been observed (PMID: 25336721), which casts doubt on the identity of the cultured species.

UQ has only been reported outside *Pseudomonadota* in one abstract, specifically in the *Bacteroidia* species *Kaistella flava* (PMID: 33724915). The quinone profile (major quinone: $MK_6$ and “a few $UQ_{10}$”) suggests a contamination likely originating from an alphaproteobacterium. The fortuitous presence of five homologs of the UQ pathway in a *Bacteroidota* genome led us to infer the presence of the UQ pathway in this phylum for the first time (S1 Table). However, the specificity of the annotations is complicated by the fact that some proteins in the quinone biosynthesis pathways belong to large families of proteins with various functions. The inference of the UQ pathway in a *Bacteroidota* genome was easily dismissed as an annotation error, since *Bacteroidota* homologs were localized in different parts of the genome rather than in compact genetic loci where *ubi* genes are usually found (Fig 3A) [1].

### S2 Text:

In several cases, the same species appears multiple times in the results (S2 Table). This can result from several situations. Firstly, the same species may have been described in different articles. Secondly, species may have been renamed, resulting in multiple entries for what is now recognized as a single species in the NCBI taxonomy. Additionally, assignments based on the GTDB taxonomy can differ from those of the NCBI, reflecting updated or alternative taxonomic criteria. Thirdly, although most of the articles mentioned only one species, there are instances where several species are mentioned in the article title, particularly when taxonomic amendments are proposed (S2C-D Figs). In such cases, the quinone descriptions found in the abstract are attributed to all species mentioned in the title.

318 species (based on the NCBI taxonomy) appear multiple times in the results. The quinone type is the same in all cases, except for *K. antarctica* and *K. jeonii.* Those two species were jointly reported in two distinct articles and were found to contain $MK_6$ in one case (PMID: 15653910) and $MK_6$ with minor quantities of $UQ_{10}$ in the other (PMID: 33724915, see S1 Text). For 254 out of 318 species (~80%), the reported chain lengths are the same. For 16 out of 318 species, the chain lengths cannot be compared because they are only specified in one abstract (S6 Table). Of the 48 remaining cases displaying differences, 42 show partial overlap (S6 Table). Most of these 48 cases correspond to species that are not the primary subject of the papers, but are mentioned in the context of taxonomic amendments, such as for *Kaistella* species (PMID: 33724915). For species (or even strains) that were described twice, for instance C*hakrabartia godavariana* (PMIDs: 31025129 and 31166165) and *Pseudarthrobacter phenanthrenivorans* (PMIDs: 31424381 and 19196765), the partial overlap in their reported quinone profiles appears to be due to the varying levels of details provided in each abstract. Some abstracts also report inconsistent information. For example, *Bhargavaea beijingensis* and *Bhargavaea ginsengi* were described as possessing either $MK_7$ (PMID: 19329597) or $MK_8$ (PMID: 22155760) (S2 and S6 Tables). These examples illustrate the rare discrepancies in our data.
