## Supplementary material for "Global distribution of isoprenoid quinones across Bacteria": S1 Dataset

>UbiX: GB_GCA_001595385.3 UbiX LVEI03000001.1_1442

Optimized sequence：

CCATGGTGGCATATCTGATTGCACTGACCGGTGCAAGCGGTGCAATTTATGGTCTGCGTCTGGCGGGTGAACTGCTGAGTCGTGGTGATGATGTTGAAGTTATTATTAGTCCGAGTGGTTTTCTGATTCTGAAAGAAGAACTGGGTCTGGAATGTGCGCCGAAAGATGCAGCAAGCAAAATTCGTGCATATCTGGAAGGTCAGGGTCGTGCGCTGAAAGGCCGTCTGGGTATTACAGCACATGATGATATGAGCGCATCTGTTGCAAGCGGTAGTAGCCTGCTGAAAGCGATGATCATTTGCCCGTGTAGCATGGGTACCCTGGCACGTGTTGCAAGCGGCGTTAGTGGTAACCTGATTGAACGTGCAGCAGATTGTATGCTGAAAGAAAAGCGCCCGCTGCTGCTGGTTCCGCGTGAAACACCTCTGAGCAGCATTCATCTGCAGAATATGCTGCGTCTGTCACGTGCAGGTGCAGTGATTCTGCCGGCAATGCCGGCATTTTATCATAAACCTTCAACCATTGATGATATGGTGGATTTTATGGCAGGTAAAATTCTGGATATGCTGGGTGTTGAAAATAGCCTGTATAAACGTTGGAAGAAGGAAGCCGAATAAAAGCTT

>UbiV: GB_GCA_001595385.3 UbiV LVEI03000001.1_1680

Optimized sequence： CATATGGAAATCACCCTGGGTCCGGTTCTGTTTGATTGGCCGAAAGATGAAGTTCTGAAATTTTATGAGGAGGCAAGTCGTATGGATGTGGATAGAGTGTATATTGGAGAGGTGGTGTGTACCCGGAAAATTGGTCTGCGTATGAATGACATTGAAGGTATAATTAAGCTGCTGCAGGATTCGGGAAAAAAAGTTATTCTGAGCACCCTGGCAGTTATTAGCAATGAAGAAGAACTGGAATTTACCCGTAAACTGCTGCATCTGCCGTGTCCGGTTGAAGCAAATGATATGAGCGTTTTTAATATGGCAGGTGAACGTGAACTGGTTGCAGGTCCGCATATTACCGCATATAATGCACCGACCATTGAATTTTTTAAAAGCATTGGTGTTAAACGTGTTGTTTTTCCGGTTGAACTGCCGAAAGCAAGCATTGAACATGATCTGCGTGCAACCGGTATTTTTGGTGAAGTTTTTGCACATGGTAAAGTTCCGCTGGCATTTAGCTGGCGTTGTTATACCAGCCGTGCATTTGGTCTGAATAAAACCAATTGTAAACATCATTGTATGAAATATCCGGATGGTATGGAACTGAAAACCGTTGATGGTGAACCGATTTTTAGCGTTAATGGTACCAGCATTCTGAGCGCAAGCACCTATAGCCTGGTTGAATTTGTTGAAGATCTGAAAGGTATTGGTGTTGGTGCACTGCGTATTAGCCCGCAGTATCGTAATACCGCAAAAATTGTTGAAGTTTTTCGTGCACGTGTTAATGGTACCCTGGGTCCGGGTGAAGGTATGAAAGAACTGAAAGCAGTTACCGAAGGTAGCTTTAGCAATGGTTGGTATCATGGTGGTGCAGGTAAAGAATATCTGAATGCAGTTCTGGGTTAAGGATCC

>UbiU: GB_GCA_001595385.3 UbiU LVEI03000003.1_59

Optimized sequence：

CATATGACCAGACCAGAAGTTATTGCACCGGCCGGCAACCTGGCAAGCCTGAAAGCAGCAGTTGACAGCGGCGCCGACGCCGTTTACCTGGGCTTTAACGACGCAACCAATGCACGCAATTTTGAAGGCCTGAATTTTACCAGCAGCGAATTAGCAGAAGGAATCAAATACGTTCGGAGCAAAGGGAGACAGTTTTATGTTGCGATCAATACCTTCCCACAGGGCGAAGACTTTCCGAAGTGGTATAAAGCAGTTGACAGTGCAATGGAGATGAAAGCAGACGCAGTTATTATTGCAAACGTGGGAGTTTTACGGTATGCACGCCAGAAGTATCCAGATGCAACACTGCATCTGAGCACGCAGGCATCAAGTTCAAATTATGAAAGCATTAACTTCTACCGTAAGCACTTTGGTATTAAAAGAGTTGTTCTGCCCAGAGTTTTAACCATTGAAGAAATTAAACACCTGAAGGAACGCACAGAAGTTGAGATTGAAGTTTTTGCACTGGGTGGTCTGTGTATTAACATTGAGGGCCGCTGTTATTTATCCTCATACGTTACCGGAGCAAGCACCAACACCGAAGGTGCCTGTAGTCCGAGCCGTTTTGTTAGATTTAATAGTAATGACGACGGTGGTATGAGCATTGCACTGAATGGTATGACCCTTAATAAACTGGGCAAAGATGAAAGCAGCCCATATCCGACCTGTTGTAAAGGACGTTATACCGGTCCCGACGGCGGCTATGCATATATTTTTGAAGAGCCGGAATCGCTGAACGTTCTGGAGATCATTCCGGGTCTGATAGATGCAGGTGTTGCCGCACTGAAGATAGAAGGTCGTCAACGTACCAAAAGCTATGTTGGTGCCATGACCAGAGTTATGCGGGAAGCTGTGGATAGCTGTTGGAAGGATAGAGCAGGTTATACCGTTAGACCGGAATGGAGCAGCAAAACAGTGGCGACCTTTGAAGGAAGCCGTCAGACCCTGGGCAGCTATCTGACAAAATAAGGATCC

>UbiT: GB_GCA_001595385.3 UbiT LVEI03000001.1_1685

Optimized sequence：

CATATGGAAGAAATCAAGAAGCAGCTGAGAGAAGAATTATATAAAGGACTGAGACTGCCGCTGAAAGCGATACCGCTGTGGATGGAAGCAATCGGGGTGGGGGTGTTTATAAAAAGCATCCTGGAAAAAAATCCGAGCTTTCGTGAACGTCTGGGTGAACTGGATGATAAAGTTTTTATGTTTGAAGCAAAAGATCTGGGTAAAGGTTTTTTTATGCATATTAAAGATAATGATATTAAAGTTAAACCGCATAGCGTTCGTGCACCGGATGTTACCATGAAAGGTGAAATGAGCGTTCTGATGGATGTTCTGCTGGGTAAAGAAGATCCGGATACCGTTTTTTTTAGCCGTAAACTGGAAATTACCGGTGATACCGCAACCGCAATTCATTTTAAAAATCTGCTGGCAGCACTGGGTTAAGGATCC

>UbiA: GB_GCA_001595385.3 UbiA LVEI03000001.1_1416

Optimized sequence：

CCATGGTGGGTCAGGTTGCAGCAGATAAACTGCATGCAGTTAGTGAACTGCTGCGTCTGCCGCGTCAGCAGGGTACACTGCTGCTGCTGTGGCCTACAATGTGGAGTCTGTTTATGGCGAGTGGTGGTCGTCCGGAACTGAAATATCTGAGCATTTTCATTATCGGTGCTTTTCTGATGCGTTCAGCAGGTTGCGCCGTTAATGATATTGCAGATCGTGATTTTGATCCTCATGTGGAACGTACACGTACCCGTCCGATTGCTTCTGGTCGTCTGAAAGTTAAAGAAGCCATGCTGGTTTTCGCCCTGCTGAGCGCAGTTGCCTTTGCCCTGGTTCTGCAGCTGAATCGTCTGACCGTTATGCTGAGCCTGGTTGCCCTGGCCCTGGCAGGTGCATATCCATTTGTTAAACGTTTTAGTCATTTTCCTCAGGTTGTTCTGGGTATGGCATTTGGTTGGGGTGCAGTTATGGCGTGGTCCGCAGTTCGTGAAGAAGTTGGTGTTGCAGCACTGCTGATTTTCACCGCGAATATTTTCTGGAGTACTGCATATGATACTATTTATGCACTGATGGATCGTGATGATGATATTAAAATTGGTGTTAAAAGTACAGCCATCTTCTTCGGTGGTTCTGTTTATAAAGCACTGAGCGTTCTGTATCTGTGTTTTGCAGTTGCCCTGGGTGCCGCCGGTATGGTTGTTGGTCTGGGTGGTATTTTCATGACCGGCCTGCTGATTTGTCTGATTCTGAGCCTGGCCATTGTTGAATTTGTTAAGAAGGAACGTACCCGTCAGGCTGCGTTTAAAGGTTTTCAGGCTAATGCTGCCATTGGTGGTGTTCTGCTGCTGTTTATTATTCTGGATATGAATCTGTAAAAGCTT

>UbiC: GB_GCA_001595385.3 UbiC LVEI03000001.1_1415

Optimized sequence：

CCATGGCTAAAGGCTTTAGCTATACCCTGCTGGGCCAGTGGCTGGGTGTTGAAGAAGCGCGTCGCAAAACAATCCTGGATGGTCTGCTGCCGCATCAGAAACTGCTGCTGTTTAGCGAAGGTAGCATGACCCTGGAACTGGAACTGCTGACCCAGGGTAATGTGGAAGCAGAAATTCGTTTTATGGGCCTGACTAGTATTACCGCCGAAGCAGCCTCATTTCTGGGTGCCGAAGTGGGTGCAGAAGCAATGGAACGTGAAGTGTGGCTGACCGGTGGCGGTCGTCGTCTGCTGTATGCACATGCACTGATTCCGGAAGGCATGATTGCACCGGATATTAAAAGTGCACTGGATGAACGTCCGAAAGAACCGCTGGGTCGTGTTCTGGCCAGTAATGGTGTTCTGTTTGCAAAAGAACGTCTGGAAATTGGTATTGTTAAAAGCCCGTGTGCAAGCCGTGATCTGGAAATTCCGGAAGATACCCCGCTGTTTGCTCGTCGTTATATTCTGTTTAATAAAGGTGCCGATCGTTGGATTATTAAAGCCGGTCTGACCGAAATTTTCAGTCCTGAACTGGTTGGTGCAGTTCTGCGTAGCTAAAAGCTT

>UbiD: GB_GCA_001595385.3 UbiD LVEI03000001.1_1417

Optimized sequence：

ccatggcgccctactatgatctaagggaatttatagaggtcctggagaagaagggtttgctgaagcgcgttaaaaccgaagttgacccggttctcgaaatcgctgctattcaagaacgtctggtgaaaagcggtggtccggcggtgttgtttgaaaaagtgaaaggccaccgcatggcggttttgggaaacctgttcggcaccgcggagcgcgttgcgctgggcctgggcgtcaccgaagaagaattatcggatattggccagttcatagcgacgctgcaacgtccgcagccgcctgagggtctgtgggatgcggtgaagaagatcccgttcttcggtaagatcctgaccctgggtccgaagacggttaaatctgcaccgtgtcaggatgtcgtggagacagacacggcggatctgtccaaaattccgatcattaaatgctggccgggtgatgctgcgccactgatcacctggccattggttgtaactcaaagcccgcaaggtggtccgtataacgtgggcgtctatcgcatgcagtacctggatggtaagcgcgcgatcatgcgttggctgtcccatcgtggtggcgcaactcatcagcggctgtgggaaaaggagggcaaagcaatgccggtggcggtggccatcgggtgcgatccggcgacgatcatcgcgggcgtgacccctgtgccggaggacgttggtgagttccactttgcaggtgtattacgtaagaaggctatcgaattagttgagtgcaaaaccattccgttgaaggtgccggcgaccgcggaaattatcattgagggtgagatccgtcacggcgaactggaaatggaaggcccgtttggtgaccataccggttattacaacgctgctgaaccgttcccggtatttcatgttaaagccatcacccaccgtaaagatgccatctatatgaccaccattaccggccgtccgccgaaagaggacgctgtgatcggcacggttttgaataagctgtaccttccgagcctgaagctgcaatttccggaagttgttgatttttgtctgccgatggaagcagtttcctaccgcattgccgtggtgtccattaaaaaggagtacccgggccacgcaagacgtattatgatgggcctgtggggtgttttgaagcagttcatgtatgttaaatacattatcgtggtggacgacgacgtggacgtccacaactggaccgatgtcatctgggcaatcagcacccgtgttgaccccaaacgtgataccttgatcattgagaacacgccgatcgattatctggacttcagcagcccgattgagaatctaggtagcaaaatgggtattgacgcgaccaataagtacccgccagaggtgagccgtaaatggggtgagaaaatggaaatggacagcaaagttgaggaactcgtagagaagaagtggaaagagtacggcttttaaaagctt

>UbiG: GB_GCA_001595385.3 UbiG LVEI03000001.1_1418

Optimized sequence：

Ccatggcgactacagagtcacagaaatttgaacaatatggaagtgattggtggaatccggcaggccgtctgttctccttacatcgtattaacccactgcgcttcggttacttctccagccgtagcggcgaactggcgggcaagaccgttttggacatcggctgcggtggtggcctgctgagcgaggagttcgccaaggcgggcgcaaccgtcaccggtattgacctgtctccggttgcgattgacgctgcgaaaggtcactgcgcggcgtctggcttgtcgatcgactaccgcgtggcctccgtggagaaaacggctcgtgaaggtaaacaatttgatgttattgtctgtgcggaagttctggagcacgttgatgacttaaatggttttcttagagacagcctgtctatgctgaagcacggcggtctgttctttttcggcaccatcaacaaaacgtttaaggctcgtttcctcgctctgtttatggcagaagacgtgttgggtatggttccgcgtggtactcatgattataaccgtttcgtgcgtccgagcaccctgaaagaaatcctagcgcagaacggggtggagatcgaagagctgaagggtatgagctatgatccgttgcgcttggagtttaagatcagcaatgataccagcgtgaactatctgggctacgcacgcaaaaaataaaagctt

>UbiB: GB_GCA_001595385.3 UbiB LVEI03000001.1_1682

Optimized sequence： ccatggcggcttcagcatataaaaatataagaaggctaaaccgtatcgttattaccctgatccgctacggctttggtggcttagcacgtgacctgcgtgtactgccgagttttgttccggcaatcgagcgattgttcatctccaagaaagcgcgtgatttatccgccccggtgcgtattcgtctagtgcttgaagagctgggtccgaccttcattaagctgggtcagatcgccagcacccgtgcagatatcctgccaccggattgggtggaggaattcaagaagctgcaagatatggttccgccggttgagttcgaggaagttcgtcgtattatcgagggcagcttcaaggcgcctattggggcgaagttcgcctcgttcgacaccgaaccggtggcgtctgcgagcatcgcgcaggttcattatgcggaactgttcgacggctccaaggtggccgttaaagttcgccgtccaggtatcgagcgcgtcattgcgtccgacattagcgtcatgcacaccattgccggtctactggatcgctacgtgagtgctgctcgtcgctaccgtccgcacgaagttgtttctgaattcgagcgcgtgatcaagtccgagcaggatttgacgattgagggcgtgaacttgaatcgtttcagcgacattttcaaagacgacccgcacatccagatcccgcgtgtgttctgggattacaccactgaggacgtgctgacgatggagcgtatttctggcaccccgatcgacgaagtcgaaactctgaaaagcaagggtattgacatcaaggaggttgcggttcgcggcattggcattttctttaagcaggtttttgagcacggcatctttcacgccgacctgcatccgggtaacattttcgtgcgtgatgatggtgttatcatttatctggattttggtatcatcggccgtctggaccgtgatctgcgtaagtatctggcgagcatgctgtttcatctggtgcgcagcgattattaccgcatggctttagtgcaccgtgaaatgggtttgatcggcgatgatgtttccctgagcgagttcgaagaagcgttgcgggacattagcgtaccgatctttggtaaatcgctggagaaaatcgacatcagcggtctgctgatgaaactcttacagaccgcgaaacgttttaacatgaaattgcaaccgaatttgctcctgttacaaaaaagcatggtgataatcgagggtgtcggtcgtcaattgtacccggacgtgaacatgtgggaagtggcaaaaccactgattttcaaatggatggctaaagaaaagctgtctccgaaaatgatcgcggaacgtggccgcgaaaaactggaggaaattatggaaacggcgtttgacttgccggtcaattttaacaccctgctgcgcagaacgctgcgcgaggacatcaagatcggcttcgtgcatcatagactcgaagaggtcaccaatgagctggagaatgcaggccgtcgtattggtggcggtatcgtggttgcggcactggtcattggtgcgtcgctggtggcagttttcagcaaagaggcgaccacgtttctgggcctcccggttttgagcggcgctggttatttggttgcagcgatcatgggtctgcgtttgtttcgtcgtagcggacgcaacggtaggtaaaagctt

>UbiE: GB_GCA_001595385.3 UbiE LVEI03000001.1_1684

Optimized sequence：

Ccatggcgtcagaaaaaatgacacactttggaaatcaaactatcccggaaggtgagaaagagaagaaggtgcgtgaagtttttgacagcgttgctagccgttacgacctgatgaatgatttgatgagcttcggtgttcaccgtttctggaaacgttttgttgcggcggaaaccggcctgcgtccgggccaaagcgcgattgatgttgccggtggaaccgcagacatcagcctgctgatggcagaccgcgtgggcgaggccggcaacatcgttgtgtttgacatcaacggtgagatgctgaagtatggtaaagagaaatgtgttgatcgcggttacttgaagaacatccgcttcgtgcagggtaatgcggaagatattgcgttcgacgacaacaccttccactgcgctaccgttggttttggcattcgcaacgtgacgcatctggatcgtgcttttcgtgagatgacccgtgtcgtgaaaccgggtggtaaagtcatctgcctggaattttcccatccgacgagcaaactgttcaaaaaggcgtacgatttatattcgttctcttttattccgaatgtaggcgagatgattaccggtaatcgctctgcgtatgaataccttccggagtccatccgtaaattcccaccgcaggaggaattgaagaagattatggaaggcgcaggtctgtggaaagtgaagtaccataacctcatgaacggcatcgccgcggttcacgtgggcgtcaaggtgtaaaagctt

>RquA: GB_GCA_001595385.3 RquA2 LVEI03000001.1_991

Optimized sequence：

ccatggacatatatgaattacaagattcaaggcccctagctgaggaacgctggctgcggaagggactgttttatcgttactttctggacggcgtgccggattacctggctcgtaactattggtgggcctacttatggaaaccgggcgcatggtttttcgaccaccagccgattatcaatgcgatcctgttcggccaataccagcgtcttatgggtgaaacgctgcgcgtgatccaagcaagacctagcggtcgtatgctgcaactgtcttgcgtttacggcaagctgaccccgagtttggcgggtctggactctcgtcgtttgcacctgaccgatgtgtccccggtgcagctgggcattagcatgcgtaaggcgggcgagaaactggtggccacccgtatgaacgcggagagcttgggttatcgtgacggtgttttcgacaccgttctcatttttttcctgatgcatgaaatgccgccagaagctcgtcgccgtactctcagcgaagccatccgcgtactgtcgccgaagggccgtttggtcatcaccgaatatggtcatgagccgaatgcgaacccgatctatcgctttcgtttgtccagatggattattggtaagttggaaccttttctgccgggtttctggcgtgaggaactggatgttagcatgaaaaacgcggcaagcattaatcgcaagacgatcaaaaggaacggtaaagatgttccggttttcaaaggcttctaccgtgtcgcggagtacgaggtggagtaaaagctt
